# Revisiting sour rot of grapevine through disease-associated microbiomes: a tripartite co-infection?

**DOI:** 10.1101/2024.09.19.613941

**Authors:** Matthieu Wilhelm, Hévin Sébastien, P Patrik Kehrli, Bart Buyck, K Katia Gindro, Jean-Luc Wolfender, Valérie Hofstetter

## Abstract

Sour rot in grapevines is thought to result from berry infection by yeasts, acetic acid bacteria and vinegar flies. To better characterize the role of each of the actors involved in sour rot expression, we conducted experiments involving the isolation of 1593 fungi and bacteria to understand the composition and dynamics of the microbiomes associated with healthy berries, diseased berries and insect vectors. As some grape clusters became symptomatic for sour rot in the absence of acetic acid bacteria, the latter might not necessarily be needed for disease expression. Similar to other yeast genera, the yeast genus *Geotrichum* is here also reported for the first time to be able to initiate sour rot in grapes, however, this finding has to be confirmed by further studies. By allowing or denying the access of insect vectors to intact or artificially wounded grapes we could emphasize that they accelerate the expression of the disease when berries are injured. Moreover, the microbial communities identified on native vinegar flies (*Drosophila spp*.) and the introduced *Drosophila suzukii* were similar and yeast dominated. This highlights the key role of these insect vectors in the transmission of microorganisms inflicting sour rot to wounded berries. Finally, our data also suggest that sour rot and grey mould can coexist in a vineyard at an advanced stage of grape decomposition, which runs counter to recent studies that emphasize the supremacy of sour rot over grey mould.

**IMPORTANCE:** This study sheds new light on the complex interactions between microbiomes, insect vectors and physical factors favoring the development of sour rot. While previous studies suggested that acetic acid bacteria were mandatory for sour rot expression our results suggest that grape sour rot could also result solely from yeast infection. Moreover, native vinegar flies and the introduced *D. suzukii* host and vectorize a similar microbial community to injured berries in vineyards and their presence accelerates the infection process and consequently the expression of sour rot.

## INTRODUCTION

In grapevine (*Vitis vinifera*) production, sour rot is a disease that poses a major threat to grape quality. When more than 20 percent of grape clusters are affected by this disease, harvest becomes unsuitable for vinification as a result of the high level of volatile acetic acidity released from rotten grapes (Asplen et al. 2015). Prevalent in vineyards worldwide (Lee et al. 2011), sour rot causes substantial economic losses (Asplen et al. 2015, Ebbenga et al. 2021, Entling & Hoffmann 2020). The disease is known to arise from a co-infection of berries by yeasts and bacteria transmitted by insects such as vinegar flies. In the progression of sour rot symptoms, yeasts facilitate the conversion of sugars into alcohol, while bacteria catalyze the oxidation of alcohol into acetic acid. Most fermentative yeasts, except for *Saccharomyces cerevisiae*, require a minimum of oxygen to perform alcoholic fermentation (Visser et al. 1990), conditions fulfilled in injured berries (Hall et al. 2018). It is assumed that in the absence of drosophilids, the injured skin of berries heals rapidly, thereby limiting the access of yeasts and bacteria to the pulp and thus maintaining an anaerobic environment unsuitable for the growth of the fermentative yeasts associated with sour rot (Barata et al. 2012a). Less than half a century ago, sour rot was considered to be the final stage of grey mould (Bisiach et al. 1982; Zerbetto 1987). However, grey mould is now considered to be a different disease (Crandall et al. 2022), although the term sour rot is still used by several authors to refer in more general terms to grape rot (Haviland et al. 2017; McFadden-Smith and Gubler 2015). In addition, Bisiach et al (1982) and Hall et al (2018) have shown that when grape clusters infected with grey mould are surrounded by sour rot infected clusters, the expression of grey mould is inhibited. Conversely, when sour rot suffering clusters are surrounded by grey mould symptoms, sour rot expression is impaired but not completely inhibited. From these findings, the authors of the two studies deduced that the two diseases compete with each other and that sour rot supersedes grey mould when the two diseases coexist in the same vineyard.

It is widely accepted that adult flies of the genus *Drosophila* (Diptera, Drosophilidae) are the most important insect vectors of sour rot. Apart from several endemic European *Drosophila* species, notably *D. melanogaster*, the Asian *D. suzukii*, was recently introduced to Europe and North America. Both mentioned species have been shown to be involved in the development of grapevine sour rot (Hall et al. 2018, Ioriatti et al. 2018). Females of both vinegar flies typically lay their eggs in injured or overripe grape berries (Sepesy et al. 2019). The hatched larvae feed on the pulp of the berries thereby making them decay and thus unfit for consumption and winemaking (Asplen et al. 2015). It is generally assumed that native *Drosophila* species are unable to lay their eggs in intact grapes and can only do so in berries whose skins have been damaged during grape ripening by birds, insects, fungal diseases and/or heavy rainfall during grape ripening. Heavy precipitation events can cause excessive water uptake leading to rapid berry growth and skin breakdown. In contrast to native *Drosophila* species, *D. suzukii* can injure and lay its eggs in healthy grape berries with intact skins due to its serrated ovipositor (Atallah et al. 2014, Entling & Hoffmann 2020, Ioriatti et al. 2015, Kienzle et al. 2020). It is generally assumed that drosophilids transmit the microorganisms associated with sour rot from infected grapes to healthy berries during the process of egg laying (Fermaud et al. 2002). Moreover, the healing of the skin of wounded berries is delayed by the activity of hatching larvae (Barata et al. 2012a). As only *D. suzukii* is able to penetrate the skin of healthy grapes, this species is regarded as a more dangerous vector of sour rot than native vinegar flies although a recent study challenges this general assumption (Entling & Hoffmann 2020). Overall, the specific role of wounds and potential insect vectors in the development of sour rot remains unclear.

Numerous studies have extensively documented the microorganisms associated with sour rot. The yeast genera identified in sour rot symptomatic berries belong primarily to the subphylum Saccharomycotina recently divided into seven classes and 12 orders (Groenewald et al. 2023). In Europe and North America, yeasts from the genera *Candida, Hanseniaspora, Pichia* (previously *Issatchenkia*), and *Saccharomyces* are predominant (Barata et al. 2008, Barata et al. 2012b, Bisiach et al. 1982, Guerzoni & Marchetti 1987, Hall et al. 2018, 2019a, Mezzasalma et al. 2017), while two additional non-yeast genera, *Magnusiomyces* (formerly *Saprochaete*) and *Colletotrichum*, are reported from China (Gao et al. 2020). In addition, the *in vitro* assay by Gao et al. (2020) suggested that fungal moulds from genera *Aspergillus, Alternaria* and *Fusarium* could spoil grapes in China and might therefore play a role in sour rot. These same studies have also identified bacteria associated with sour rot diseased berries, predominantly acetic acid bacteria (AAB) from the genera *Acetobacter* and *Gluconobacter* in Europe and North America, as well as *Firmicutes, Cronobacter*, and *Serratia* in China. While the most abundant microorganisms in grape berries affected by sour rot are relatively well-documented (Bokulich et al. 2014, Kántor et al. 2015, Martini et al. 2009, Martins et al. 2014, Setati et al. 2012, 2015), only one study investigated, to our knowledge, the microbial changes associated with the development of sour rot (Hall et al. 2019a) and no study so far has analyzed the microbial community present on *Drosophila* adults that might consequently be transmitted to healthy grapes.

The authors report here on three experiments that were conducted to better understand the respective roles of the microbial communities, potential vectors and physical injuries in the development of sour rot in grapes. The first experiment aimed to differentiate the microorganisms associated with grey mould from those associated with sour rot in order to verify the results of previous studies that have shown that the two diseases are distinct (Crandall et al. 2022) and that sour rot takes over from grey mould when both diseases are present within a vineyard (Bisiach et al. 1982; Hall et al. 2018). To do this, we sampled net-protected bunches symptomatic of grey mould and unprotected bunches symptomatic of sour rot in the same vineyard plot. Additionally, we assessed whether the microbial communities associated with sour rot were similar across different regions in Switzerland. Findings are discussed and compared with those from other countries. The second experiment focused on the evolution of the microbiome on and in grape berries that were either intact or artificially wounded over the last three weeks preceding the harvest, thereby using insect-proof nets to regulate the access of potential insect vectors to the grape clusters. This bifactorial experimental design was adopted to elucidate the individual role of each factor associated with sour rot and to unravel the temporal dynamics of microbiome development.. Finally, the third experiment analyzed and compared the fungal communities present on the surface of healthy berries, in surface-sterilized healthy berries, in sour rot affected grapes, and on the surface of *Drosophila* flies that were captured in the same vineyard. The aim of this third experiment was to determine the respective contributions of healthy berry surfaces and insect vectors to the mycobiome associated with sour rot.

## MATERIALS AND METHODS

### Experiment 1: Fungi associated with grapes symptomatic for sour rot or grey mould

An initial experiment was conducted to compare the fungal and bacterial species present in grape clusters exhibiting symptoms of sour rot or grey mould and to assess whether the microbial community associated with sour rot varied across different regions. Grape clusters were selected based on the known symptoms of each disease. In the case of grey mould, grape clusters were considered to be infected when berries were brown and covered with the greyish mycelium of *Botrytis cinerea* (Viret & Gindro 2024). For the recognition of sour rot, skin discoloration, pulp discharge, a smell of vinegar and a minimum acetic acid content of 0.83 g/L in diseased berries were the required symptoms following Hall et al. (2018). To confirm that acetic acid concentration was over this threshold, symptomatic grape berries were crushed, and the acetic acid content of the resulting juice was analyzed using an A25 spectrophotometric autoanalyzer (Bio System, Barcelona, Spain), together with the commercial kit “Acide Acétique” Enzytec ™ Liquid (R-Biopharm, Darmstadt, Germany).

Sour rot symptomatic grapes were sampled from two vineyard areas in Switzerland, namely Nyon (Vaud) and Landquart (Grisons). In early October 2019, 18 sour rot symptomatic grape clusters were sampled from seven cultivars: Pinot noir (3 from Landquart, 2 from Nyon), Diolinoir (2 from Landquart), Pinot blanc (2 from Landquart, 2 from Nyon), Pinot gris (1 from Landquart), Sauvignon blanc (2 from Landquart), Chardonnay (2 from Nyon), and Chasselas (3 from Nyon). In addition, three clusters of Chasselas infected with grey mould in Nyon were sampled to compare them with sour rot symptomatic grape clusters from the same vineyard plot. Grape clusters were harvested using pruning shears and immediately placed in individual sterile sealed plastic bags. The samples were kept at room temperature in plastic bags for two weeks for microorganisms’ enrichment before being crushed. Subsequently, 1 ml of the material was spread onto Petri dishes (9 cm Ø) containing potato dextrose agar (PDA [potato infusion 4 g/L, D(+)-glucose 20 g/L, agara agar 15 g/L] Merk, Darmstadt, Germany). The pH of the PDA medium ranged between 5.4 -5.8, which is adapted to AAB cultivation according to de Ley et al. (1984) who found that the optimal pH to cultivate AAB is 5 to 6, although they can still grow well at pHs below 4. Petri dishes were then sealed with Parafilm®M, which is permeable to oxygen, and kept at room temperature (23-25 °C) for three weeks. We examined dishes daily over four weeks and as soon as fungi or bacteria became visible, they were isolated onto new PDA Petri dishes (6 cm Ø) to obtain them in pure culture.

### Experiment 2: Evolution of the microbial community on grapes in respect of wounds and vectors

To assess the impact of fungi, bacteria, insect vectors and wounds on the evolution of the microbial communities of berries and the expression of sour rot, an exclusion experiment was conducted in 2019. The evolution of the microbial community was monitored over the course of three weeks on healthy and artificially wounded grape clusters in the presence or absence (protected by insect-proof nets, 1 mm mesh allowing direct exchange of air and steam) of *Drosophila* flies. In a Chasselas vineyard of Agroscope at Nyon, 36 grape clusters were selected for the experiment. In early September, 18 grape clusters were protected from insects with iron cages covered with insect-proof nets, while the remaining 18 grape clusters were left unprotected. In each of these two groups, ten berries on half (9) of the grape clusters were artificially wounded using a scalpel sterilized with 70% ethanol, The experiment consisted thus of four different treatments repeated on nine independent grape clusters: protected and unwounded, protected and wounded, unprotected and unwounded as well as unprotected and wounded clusters. Grape clusters were sampled at a weekly interval for three weeks. The first sampling was carried out one week after the set-up of the experiment and consisted in pruning three clusters per modality which were then placed individually in sterile, airtight plastic bags. Back in the laboratory, two artificially wounded berries and two unwounded berries adjacent to the wounded berries were collected from each of the six wounded grape clusters. Similarly, two berries were randomly picked from each of the six unwounded grape clusters. This sampling procedure was repeated in week two and three following the wounding of the berries. Berries were brought back to the laboratory as soon as they were collected and each sampled berry opened using a sterile scalpel and twezeers Brussels used to rub berry material onto PDA Petri dishes (9 cm Ø) for 1 minute.. As in the survey described above, plates were controlled daily for the development of microorganisms. For four weeks, emerging fungi and bacteria were transferred to new PDA Petri dishes (6 cm Ø) as soon as they became visible to obtain them in pure culture.

To capture *Drosophila* flies, insect traps were positioned around the experimental Chasselas vineyard of Agroscope. These homemade traps were made of a plexiglass cylinders (20 cm in height and 10 cm Ø) in which a vinegar fly attractant was placed (Gasser lure®, RIGA AG - Paul Gasser, Ellikon an der Thur, Switzerland). The top of the cylinder was sealed with an insect-proof net, while small insects could enter through ten holes of circa 3 mm diameter drilled around the cylinder about 3 cm below the top. Traps were sampled once a week and dead drosophilids were identified under a stereo microscope. Specimens of the introduced *D. suzukii* could easily be distinguished from native *Drosophila* species by the black spot on the wings of males and the characteristic strongly serrated ovipositor of females.

### Experiment 3: Comparison of the fungal communities associated with sour rot on healthy and symptomatic berries as well as drosophilids

To determine if the fungi present on the surface of healthy berries or on the bodies of *Drosophila* species may act as potential reservoirs for the mycobiome associated with sour rot symptomatic berries, a vineyard of the Gamay cultivar affected by this disease was surveyed in Begnins near Nyon (Switzerland) in 2020. A total of 18 grape clusters (eight symptomatic and ten asymptomatic) were collected in airtight sterile plastic bags between September 14^th^ and September 23^rd^, 2020. To isolate only the microorganism community present inside of the berries, the eight symptomatic grape clusters were rinsed in tap water for one hour immediately upon arrival in the laboratory. Following this step, eight sour rot affected berries per grape cluster were sampled, rinsed several times with sterile water and left half an hour to dry under a laminar flow. We could not apply a 70% ethanol sterilization method because sour rot symptomatic berries had a damaged skin, with openings that would have let the ethanol to enter the berries and kill part of the microorganisms living inside them. Seven small pieces of each berry were sampled using a sterile scalpel and placed equidistantly on a PDA Petri dish (9 cm Ø). From the ten healthy grape clusters, eight berries were randomly collected per cluster. Half of these berries were rinsed under tap water for half an hour, then their surface was sterilized in 70% ethanol for 1 minute. Once their surface was dry, the berries were cut into small pieces with a sterile scalpel under a laminar flow. Seven samples of each of these berries (skin and pulp) were placed on PDA Petri dish (9 cm Ø). The other half of the berries were not surface sterilized and carefully rubbed individually on PDA Petri dishes using tweezers Brussels during 1 min to isolate the microorganisms present on the surface of healthy berries. As before, and for each treatment (sour rot symptomatic berries, asymptomatic surface-sterilized berries, and asymptomatic non-surface-sterilized berries), fungi were isolated in pure culture. To capture *Drosophila* adults alive, three homemade insect traps were placed in the Gamay vineyard at Begnins, with a second cylinder added inside. The bottom of this second cylinder was removed and fitted into the cylinder that contained the fly attractant with an insect net separating the two cylinders to prevent specimens from drowning in the attractant. The traps were checked daily from September 8^th^ to 14^th^ 2020. After identification of the vinegar flies, five individuals were placed equidistantly on a PDA Petri dish (9 cm Ø), with *D. suzukii* and endemic species placed in separate dishes (57 individuals for D. suzukii and 52 for endemic drosophilid spp.). The vinegar flies were pressed into the culture medium to immobilize them. As in the previous experiment, the Petri dishes were surveyed daily for four weeks to isolate the emerging fungi and bacteria in pure culture as soon as they were visible.

### Molecular characterization and identification of the fungal and bacterial strains

As soon as the fungi isolated in pure culture were sufficiently developed (0.5 cm^2^), direct amplification of the ribosomal nuclear DNA internal transcribed spacers 1 and 2 plus the 5.8S (ITS) was performed according to Hofstetter et al. (2012). For the bacteria isolated in pure culture, we amplified the ribosomal nuclear small subunit (16S) using two different primer pairs to amplify this locus in order to maximize the number of successful PCRs: 8F/1492R designed by Galkiewicz & Kellogg (2008) and 341f/785r designed by Thijs *et al*. (2017). Direct PCR was performed using a sterile pipettor tip (10 μl) to aseptically transfer a tiny amount of mycelium or bacteria in a PCR tube and to squash it manually with the tip in the PCR mix (25 μl mix, reagents, and conditions of the Taq PCR core kit (QIAGEN, Inc., Valencia, California, USA).

While it was easy to discriminate bacteria from fungi growing in a mycelial form, it was not possible to discriminate yeasts, particularly abundant in our experiments, from bacteria by visual inspection of the colonies. We therefore adopted a strategy based on the success or failure of direct amplification of the fungal ITS using primers ITS1F and ITS4, then ITS1-ITS4 alternately for isolates for which no PCR product had been obtained with ITS1F, to discriminate fungi from bacteria. For isolates for which no PCR product had been obtained with the fungal primers, we used the bacterial primer pairs (8F/1492R and 341f/785r). When direct amplification of fungal ITS and bacterial 16S had failed, we performed genomic DNA extractions according to Hofstetter et al. (2012) using material sampled from pure each isolate (50-70 mg) as soon as they were sufficiently developed (1- 3 weeks after cultivation), which were placed individually in Eppendorf tubes containing 500 µl CTAB buffer (1x). Direct PCR was performed immediately after these samples were taken. PCR products were sequenced in both directions by ^©^Eurofins Genomics (Ebersberg, Germany) or by Fasteris Life Science Genesupport (Geneva, Switzerland) using the same primer pair as for amplification.

The obtained sequences were assembled using the Sequencher v. 4.9 software (Gene Codes Corp., USA). Once assembled, they were verified by eye and sequences were then imported in MacClade v. 4 (Maddison & Maddison 2005) and for each locus (ITS and 16S) aligned manually. Since the primers used to amplify ITS are designed at the end of the small nucleic ribosomal subunit (SSU) and at the beginning of the large ribosomal subunit (LSU) for ITS1(or ITS1F) and ITS4 respectively, sequence alignment allowed ITS delimitation. The sequences of each locus were then subjected to a similarity search in Sequencer with an assembly parameter of 100% to determine the number of genotypes present in our samplings. Two different nucleotide similarity searches (https://blast.ncbi.nlm.nih.gov/) were performed for each genotype in the GenBank (National Center of Biotechnology Information). The first sequence similarity search used the “blastn” options (Megablast) and excluded “uncultured/environmental” sequences. The second was conducted using the option “Sequences from type material”. The name associated with the most similar sequence(s) (“BLAST top score(s)” expressed as %) in GenBank was adopted to name our isolates, favoring, when possible, the names associated with sequences from type material, this when BLAST top score of type sequence(s) exhibited a similarity percentage equal to the Megablast top score sequence(s). The similarity threshold values of BLAST top score(s) adopted to name sequences at systematic ranks followed Hofstetter et al. (2019) but using higher sequence similarity thresholds for species ranks. Species name associated to the BLAST top score(s) of our ITS or 16S sequences was only adopted when similarity with a GenBank sequence was 100%. When the isolated fungal species belonged to Ascomycota species complexes in which several cryptic species share identical ITS sequences, we retained only the genus rank. The suffixes cf. (likely that species) and aff. (close but might not be that species) were used for 99.5-99.9% and for 99.0-99.49% sequence similarity intervals respectively. Below 99% of sequence similarity, the isolates were assigned to genus, family or higher ranks. When assigning family and order ranks to our taxa, the classification presented in GenBank was followed. For species names we followed GenBank, but we verified for the species current name in Mycobank.

### Statistical analyses

Exploratory analyses of the data obtained in Experiment 3 were carried out using R (R Core team 2024) and the metacoder package (Foster et al. 2017). For each taxon and each pair, we performed a pairwise Mann-Whitney test (Mann and Whitney, 1947). This tests non-parametrically (i.e. without making any distributional assumption such as normality) whether the *location parameters* of two distributions are equal against the alternative that they are different (see the reference of the R function wilcox.test). We showed only the taxa whose proportions were different at the significance level of 5 % across the modalities according to the Mann-Whitney test. We controlled for the false discovery rate using the [Benjamini-Hochberg](https://doi.org/10.1111/j.2517-6161.1995.tb02031.x) (1995) procedure. However, these *p*−values should be interpreted with care given the inherent uncertainty of the test, the preprocessing steps that might affect its validity, the small sample sizes of the data set at hand and its nonparametric nature (nonparametric tests are inherently less powerful than parametric ones in order to to avoid distributional assumptions).

## RESULTS

The dataset is referenced on [Zenodo](https://doi.org/10.5281/zenodo.14958216) to ensure findability, complying with the FAIR data principles. All analyses and graphs produced with R as well as the original data are available online on a GitHub page. This allows one to fully reproduce the results presented in this work.

### Experiment 1: Fungi associated with grapes symptomatic for sour rot and grey mould

Having sampled grape clusters symptomatic for sour rot (i.e., acetic acid concentration in berries ≥ 0.83 g/L) in two regions (Vaud and Grisons) and for grey mould in one region of Switzerland (Vaud), a total of 78 fungi and bacteria were isolated in pure culture. The barcode sequences obtained from these isolates (ITS for fungi and the 16S for bacteria) corresponded to 35 different genotypes, where 11 could be assigned to the species rank and 25 to the genus rank based on GenBank BLAST top score results (Table S1, https://github.com/agroscope-ch/sour-rot/ in the “Data” folder; click on “…” in the top right-hand corner to download the files). Out of these 35 genotypes, 31 were fungi that were assigned to 12 different genera based on ITS sequences ranging from a minimum of 300 (ITS1-5.8S for 8 fungal isolates) to full ITS sequences. The four remaining genotypes were bacteria that were assigned to two distinct genera based on the 16S sequences obtained (967-1379 base pairs) using the primers 8F/1492R (Galkiewicz & Kellogg, 2008), found to amplify all the bacteria of our sampling after DNA extraction. Considering the region and disease symptoms, 12 out of these genotypes were obtained from sour rot symptomatic grapes in the Grisons, 16 from sour rot symptomatic berries in Vaud and 12 from grey mould symptomatic berries in Vaud (Table S1).

As the ITS and 16S sequences did not provide sufficient resolution to identify most of the fungal strains and half of the bacterial strains at the species level (Table S1), we inferred the composition of the microorganism communities based on the genus rank (Figure 1; in the “Figures” folder, click on Figure1.html and then on “…” in the top right-hand corner to download the file. Figures 1-7 with values can be obtained this way).

**Figure 1.**
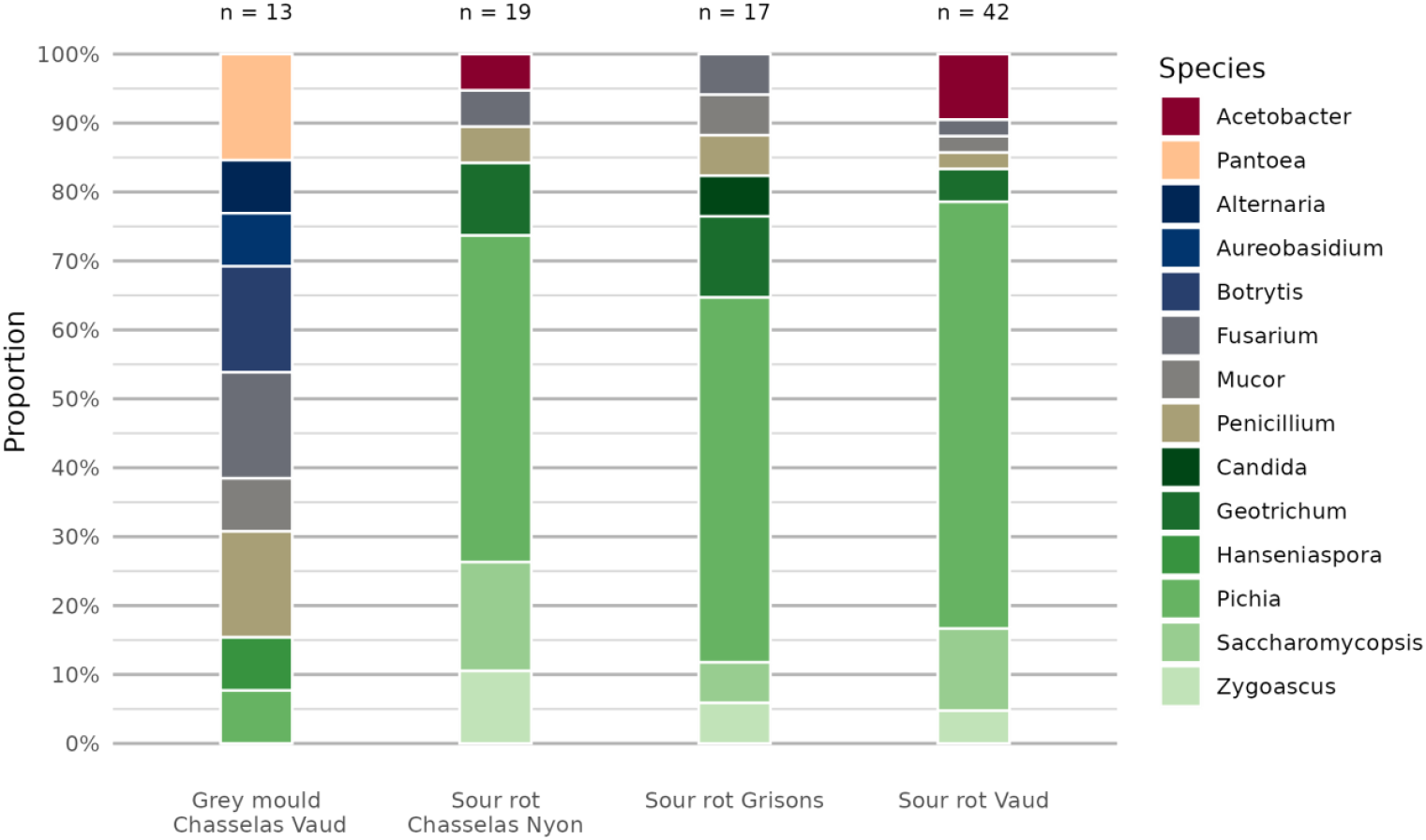
Relative abundances (expressed in %) of fungal and bacterial genera associated with grape clusters associated with: the cultivar Chasselas with net-protected clusters symptomatic for grey mould in Nyon; with unprotected clusters symptomatic for sour rot in Nyon; several cultivars symptomatic for sour rot in the two regions Grisons; with several cultivars symptomatic for sour rot in Vaud. Generic names followed the name(s) associated to BLAST top score(s) sequence(s) in GenBank (see Table S1).

By comparing the microbial communities associated with grey mould and sour rot at the genus rank in a single Chasselas plot in Nyon, significant differences were observed (Table S1; Figure 1). Only two out of the 14 genera isolated were present in both grey mould and sour rot symptomatic berries, namely the fungal genus *Penicillium* (relative abundance of 15.4% for grey mould and of 5.3% for sour rot) and *Pichia* (relative abundance of 7.7% and 47.7%). The genera *Geotrichum, Saccharomycopsis* and *Zygoascus* (all Saccharomycotina according to GenBank) were exclusively isolated from sour rot symptomatic grape clusters with relative abundances of 10.5%, 15.8%, 10.5% respectively. Together, these three genera, accounted for 84.5% of the microbial community of sour rot symptomatic clusters. In contrast, in the microbial community associated with grey mould, Saccharomycotina accounted only for 15.4% and was represented by the genera *Hanseniaspora* (7.7%), absent in sour rot symptomatic clusters, and *Pichia* (7.7%). The fungal genera *Alternaria* (with a relative abundance of 7.7%), *Aureobasidium* (7.7%), *Botrytis* (15.4%), *Mucor* (7.7%) and *Fusarium* (15.4%) were exclusively found in grey mould symptomatic grape clusters. Of the two bacterial genera identified, the genus *Acetobacter* (with a relative abundance of 5.3%) was isolated only from grapes symptomatic for sour rot, whereas *Pantoea* (15.4%) was exclusively isolated from grey mould symptomatic berries.

When comparing the microbial communities associated with sour rot symptomatic berries between the two cantons, Grisons and Vaud, the six fungal genera (*Fusarium, Geotrichum, Penicillium, Pichia, Saccharomycopsis* and *Zygoascus*) were present in both regions, while genus *Candida* was only present in Grisons (Figure 1). The Saccharomycotina yeasts accounted for 82.4% of the microbial community associated with sour rot symptomatic berries in the Grisons and for 83.3% in Vaud (Table S1). The remaining fungal genera, namely *Penicillium, Fusarium* and *Mucor*, represented 17.6% in the Grisons and 2.3% in Vaud. Overall, the fungal communities isolated from both regions were highly similar at the genus rank. However, looking at the level of species and genotypes, the two regions appeared less similar (Table S1). In fact, the two regions shared only four out of the 31 fungal genotypes, namely *Pichia californica, Fusarium* sp. 4, *Penicillium* sp. 1 and *Zygoascus meyerae*. Species of *Pichia* differed between regions, with *P. kluyveri* and *P. membranifaciens* (with two isolates cf. *membranifaciens*) only isolated in the Grisons, and *P*. aff. *manshurica, P*. aff. *kluyveri*, and *P*. cf. *kudriavzevii* only isolated in Vaud. Two different species of *Geotrichum* were isolated in each region, while different genotypes of *Saccharomycopsis crataegensis* and of *Mucor* cf. *circinelloides* were present in each of the two regions. Looking at the bacteria, only the genus *Acetobacter* was isolated in Vaud with a relative abundance of 9.5%. while no bacteria were isolated in the Grisons (Figure 1).

### Experiment 2: Evolution of the microbiome on grapes in respect of wounds and vectors

The aim of this experiment was to follow the evolution of fungal and bacterial communities preceding the expression of sour rot and to determine the contribution of wounds and insect vectors on disease expression. Berries were sampled from grape clusters treated under four different modalities: undamaged berries protected from insect vectors by a net, protected by a net and artificially wounded, unprotected and undamaged, and unprotected and artificially wounded. Sampling was conducted one, two, and three weeks after the wounding of grape berries. Trap captures confirmed that potential insect vectors such as *Drosophila* spp. and *D. suzukii* were present over the experimental period.

A total of 495 microorganisms (fungi and bacteria) were obtained in pure culture after sampling these four modalities (Table S2, https://github.com/agroscope-ch/sour-rot/ in the “Data” folder; click on “…” in the top right-hand corner to download the file). In total, we obtained 469 fungal ITS sequences representative of all the cultured morphotypes, except one (anamorphic mycelium named fungal sp.), whereas a total of 25 sequences for the 16S locus were obtained for bacterial strains. Overall, 90 different sequence genotypes were obtained, 81 ITS for fungi and nine 16S for bacteria. BLAST top score(s) results (Table S2) assigned the ITS genotypes to 34 fungal genera and the 16S genotypes to three bacterial genera. Out of the 81 ITS fungal genotypes, only 28 allowed an identification at the species rank, while merely two out of the nine bacterial 16S genotypes could be assigned at species rank.

As sour rot has been shown to result from a tripartite co-infection involving fungi, bacteria and insect vectors, we looked at the evolution of the microbial community over time at genus/species rank (Table S2) considering only the wounded and unprotected berries as these are susceptible to express sour rot symptoms (Figure 2). Among these grape clusters, only the three clusters sampled after three weeks were symptomstic of sour rot (acetic acid content = 1.4, 2.2 and 4.6 g/L). Detection of sour rot associated fungi started after three weeks, with Saccharomycotina representing 70% of the microbial community, while the AAB reported to be associated with sour rot were still absent, with only representatives of *Bacillus* were isolated (Table S2). Representatives of four yeast or yeast-like genera were isolated, namely one *Zygoascus* species (*Z. meyerae*; Table S2) accounting for 15% of the microbial community, one *Saccharomycopsis* sp. (a cf. sp. likely to be *S. vini*) with a relative abundance of 20% and two unidentified species of *Geotrichum* (sp. 1 and 2) representing 35% of the microbial community.

**Figure 2.**
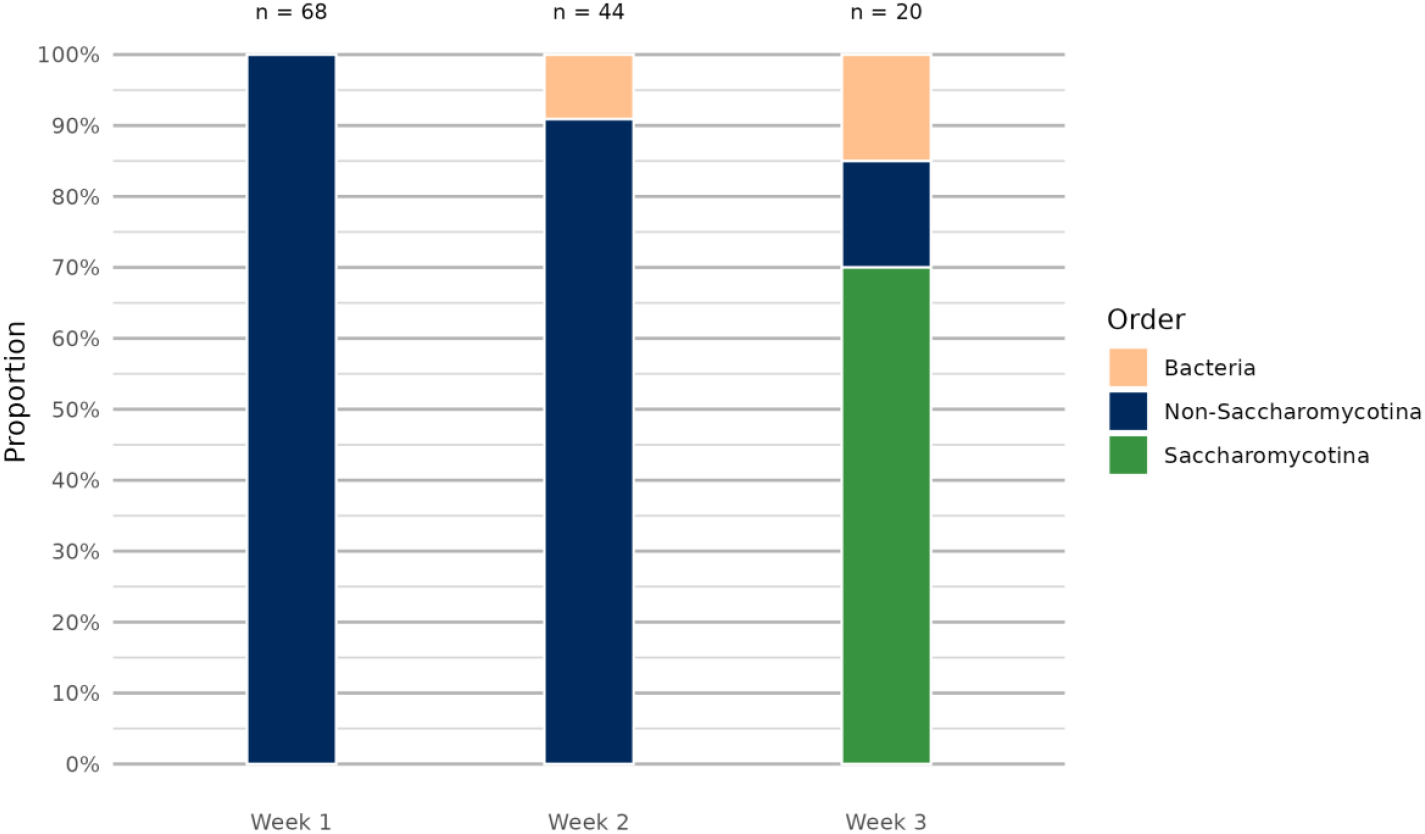
Evolution of the relative abundance (expressed in %) of sour rot associated microorganisms in wounded and unprotected grape berries of Chasselas over a three week period after their wounding. Sour rot involved yeasts (or yeast-like fungi) are indicated, while all fungi that are not responsible for sour rot expression are grouped as “Non-Saccharomycotina” (see Table S2 for the species/genera names assigned to our sequences based on BLAST results).

To determine whether sour rot affected berries transmit disease-causing microorganisms to adjacent unwounded berries in the presence of insect vectors such as drosophilids, we compared the identified microbial communities associated with wounded berries affected by sour rot to the microbial community on unwounded berries adjacent to symptomatic berries as well as with healthy berries sampled from unwounded grape clusters three weeks after the starting of the experimentation in unprotected grape clusters (Figure 3). The microbial community structure of symptomatic berries differed strongly from that of unwounded berries. This was especially evident for Saccharomycotina, which represented 70.0% (Dipodascales 50.0% and Ascoideales 20%) of the microbial community in sour rot affected berries, but only 14.3% (Dipodascales) and 16.7% (Ascoideales) for those associated with adjacent and healthy berries, respectively. In addition, two other orders of fungi were present in the three different set-ups, namely Hypocreales (symptomatic: 10.0 %, adjacent: 23.8% and healthy: 25.0%) and Agaricales (5.0%, 9.5% and 25.0% respectively). Unwounded berries, adjacent to wounded berries or taken from healthy clusters, shared the fungal order Eurotiales (19.0% and 8.3% respectively), an order absent in symptomatic grapes. The fungal orders Polyporales (9.5%), Trichosphaeriales (19.0%), and Mucorales (4.8%) were present only on berries adjacent to injured berries and the order Pleosporales only found on berries from healthy clusters (25.0%). Considering the bacterial community, the order of Bacillales was only present in symptomatic berries with an abundance of 15.0%. Overall, the microbial communities isolated from unwouded healthy berries were more similar to each other than to the community isolated from symptomatic berries because of the dominance of Saccharomycotina and the presence of bacteria (Bacillales), already present the second week, in sour rot symptomatic berries.

**Figure 3.**
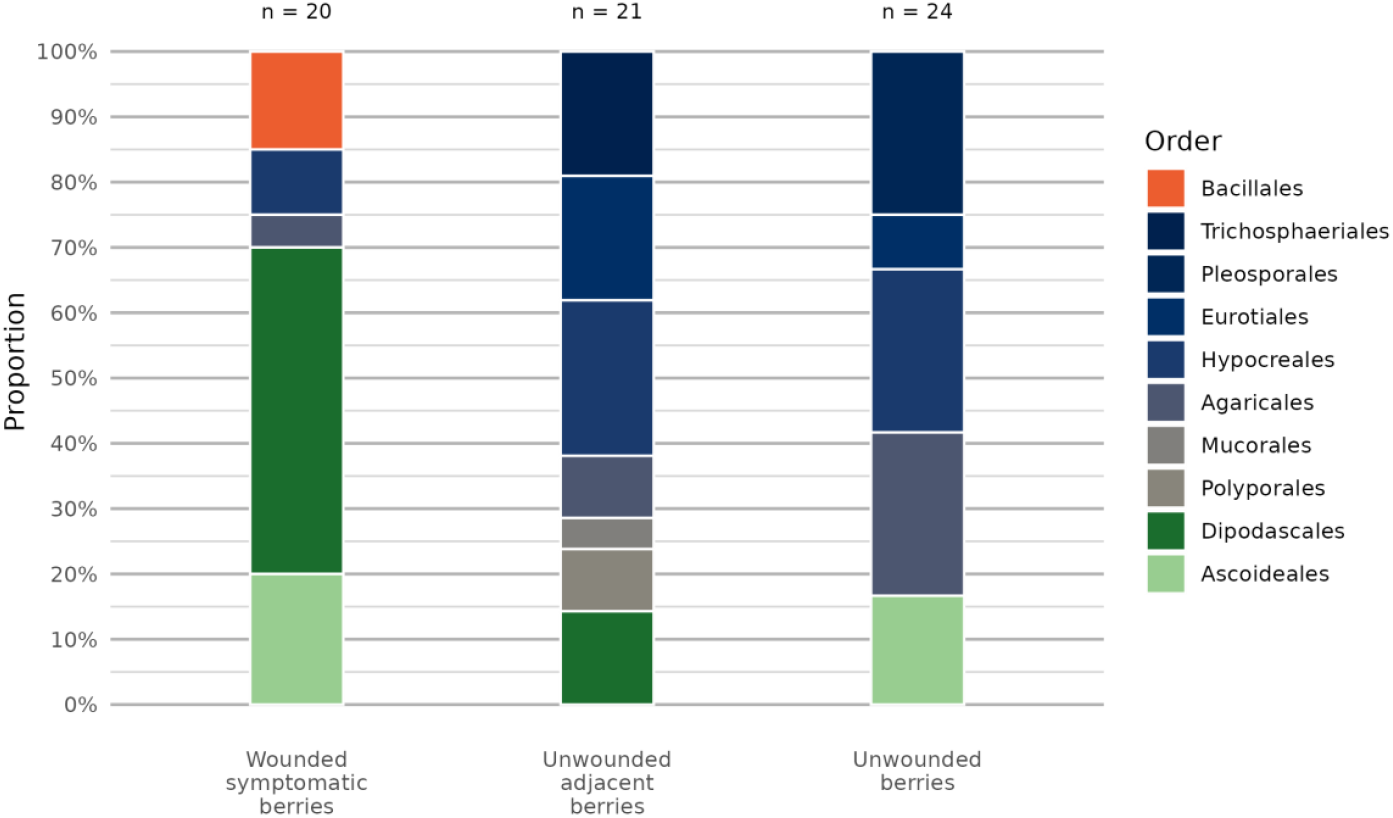
Comparison of the relative abundance of microorganisms isolated from unprotected grape clusters of Chasselas three weeks after the start of the experiment (see Table S2 for the classification assigned to our sequences).

We also tested for the impact of insect vectors on the development of sour rot in the absence of wounds, comparing the evolution of the microbial communities on unwounded healthy berries protected or not by a net over the three weeks (Figure 4). Although insect vectors such as *Drosophila* spp. could access grape unprotected grape clusters, none of the unwounded berries developed symptoms of sour rot over the three-week exposure period. One week after the installation of the protective nets, the microbial community was still the same as for unprotected berries and dominated by Hypocreales (Figure 4). In addition to Hypocreales, the two fungal orders Xylariales and Eurotiales (both orders at 7.1% relative abundance) were also present on the surface of protected and unwounded berries, whereas the bacteria belonging to Lysobacterales (3.3%) were only present on the surface of unprotected berries. After two weeks, Hypocreales on unprotected berries strongly decreased and the order were largely replaced by fungi belonging to Agaricales (37.1%), Microascales (20.0%), Eurotiales (8.6%) and Trichosphaeriales (5.7%), while the abundance of the bacteria (Lysobacterales) slightly decreased (2.9%). On the surface of net-protected berries, however, Hypocreales remained dominant (66.7%) but were partly replaced by Eurotiales (20.8%) and Agaricales (12.5%), while the order of Xylariales disappeared completely. After three weeks, species of Pleosporales and Saccharomycotina appeared, particularly on the surface of unprotected berries (25.0% and 16.7%, respectively), while the Xylariales (23.8%) reappeared and the bacterial order Enterobacterales (4.8%) was identified for the first time on net protected berries. Although the microbial communities evolved over time, they were relatively similar between the two setups after three weeks (Figure 4). They both included the four fungal orders Agaricales, Eurotiales, Hypocreales, Pleosporales and each one order of Saccharomycotina (Ascoideales on unprotected berries and Saccharomycodales on protected berries), with the additional fungal Xylariales and bacterial Enterobacterales in the net-protected modality.

**Figure 4.**
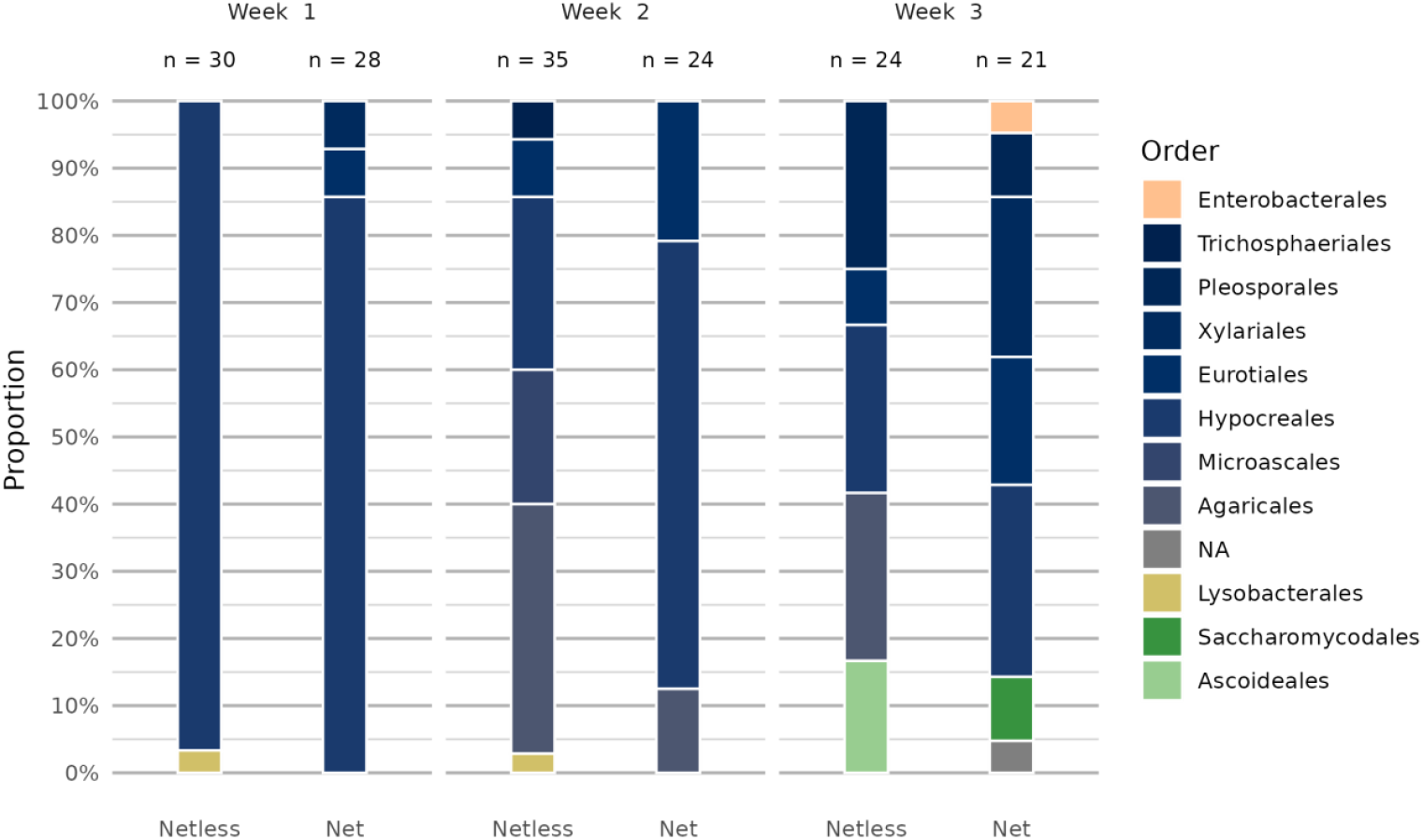
Three-week evolution of the microbial community inferred from the relative abundance (expressed in %) of fungal and bacterial orders present on the surface of unwounded berries from grape clusters of cv. Chasselas protected or not by a net in the vineyard plot of Nyon (see Table S2 for the classification of isolates at order rank).

Interested in the effect of wounding on the evolution of the microbial community, we compared the evolution of the microbial communities associated with wounded and healthy berries, all sampled from unprotected grape clusters over the three weeks before harvest (Figure 5). Already one week after wounding, the microbial communities on unwounded and wounded berries differed considerably. While the microbial community on the surface of unwounded berries was entirely composed of Hypocreales (96.7 %) and Lysobacterales (3.3%), the microbial community on the wounded berries was more diverse. Hypocreales still dominated the community (57.3%), followed by Agaricales (27.9%), Microascales (8.8.0%), Xylariales (4.4%) and Eurotiales (1.4%). After two weeks, the microbial community on the surface of intact berries closely resembled that of wounded berries after one week, with the same four most abundant orders representing more than 90% of the microbiome. However, on wounded berries the fungal order Pleosporales appeared with an incidence of 9.1%, while the Microascales disappeared. By week three, the community on the surface of unwounded berries closely resembled that of wounded berries after two weeks, being composed of the four fungal orders Hypocreales (25.0%), Eurotiales (8.3%), Agaricales (25.0%) and Pleosporales (25.0%) and Saccharomycotina (order Ascoideales, 16.7%). But two Saccharomycotina orders, Ascoideales (20%) and Dipodascales (50%), largely dominated the microbial community associated with wounded berries after three weeks (70.0%) and were accompanied by the two fungal orders Hypocreales and Agaricales. Yet, only the bacterial order Bacillales was isolated from wounded berries in the third week with a relative abundance of 15.0%. No acetic acid bacteria (AAB) were isolated even though three grape clusters became sour rot symptomatic according to their acetic acid content (≥ 0.83 g/L).

**Figure 5.**
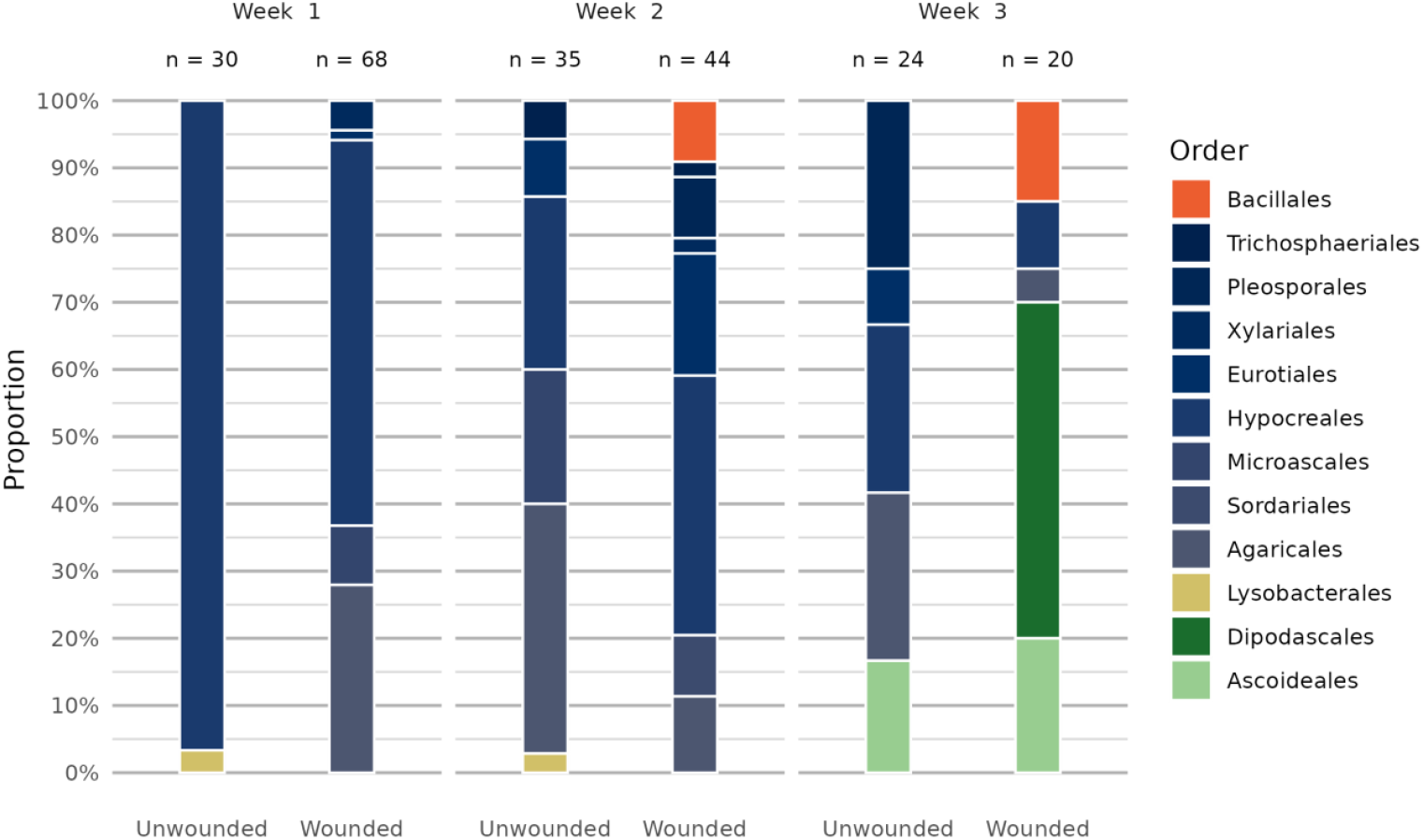
Three-week evolution of the microbial community inferred from the relative abundance of fungi and bacteria present in berries of Chasselas sampled from unprotected grape clusters of artificially wounded or intact berries in the vineyard plot of Nyon (see Table S2 for the classification of the isolates).

We also tested for the impact of insect vectors on wounded berries. For this, we compared the evolution of the microbiomes associated with wounded berries protected or not by a net over the three weeks (Table S2, colums A-F). Over the first two weeks, the evolution of the microbial communities on wounded berries in the presence or absence of potential insect vectors was similar in respect of the fungal orders present, although there existed some difference in their relative abundances (Figure 6A; Table S2). Hypocreales (genera *Fusarium* and *Sarocladium*) dominated both modalities after one week, followed by Agaricales (*Coprinopsis*). Together, these three genera accounted for 77.0% of the microbial community isolated from net-protected wounded berries, and for 85.3% of that from unprotected wounded berries. After the second week, the two communities shared four more fungal orders (Eurotiales, Xylariales, Trichosphaeriales and Sordariales) as well as the bacterial order Bacillales. Together all seven orders represented 85.2% and 90.9% of the microbial communities isolated from wounded berries protected with a net or unprotected, respectively. However, at the genus rank, the microbiomes of the two modalities shared only the five fungal genera *Aspergillus* (Eurotiales), *Apiospora* (Xylariales), *Sarocladium* (Hypocreales), *Coprinopsis* (Agaricales) and *Gibellulopsis* (Trichosphaeriales) as well as the bacterial genus *Bacillus*. These six genera accounted for 73.3% of the microbiome on net-protected berries and for 54.6% on unprotected berries. Three weeks after the wounding of berries, the communities of the two modalities were not only more diverse but clearly different among the two set-ups sharing only three out of the 26 isolated genera. The order Pleosporales was represented by six genera (*Alternaria, Preussia, Pithomyces, Epicoccum, Pseudopithomyces* and *Torula*) and totalled 37.7% of the microbiome on net-protected berries, whereas it was entirely missing on unprotected berries. With a relative abundance of 70%, the Saccharomycotina (Ascoideales and Dipodascales) represented by the genera *Saccharomycopsis*, and *Geotrichum* and *Zygoascus* respectively, was the most abundant group in the unprotected modality, while in the net-protected berries these same three genera were absent. Another yeast order (Saccharomycodales) was present on net-protected berries, the genus *Hanseniaspora* (4.9%). In the third week, Bacillales (genus *Bacillus*) was only present in wounded and unprotected berries, with a slightly lower relative abundance than after two weeks.

**Figure 6.**
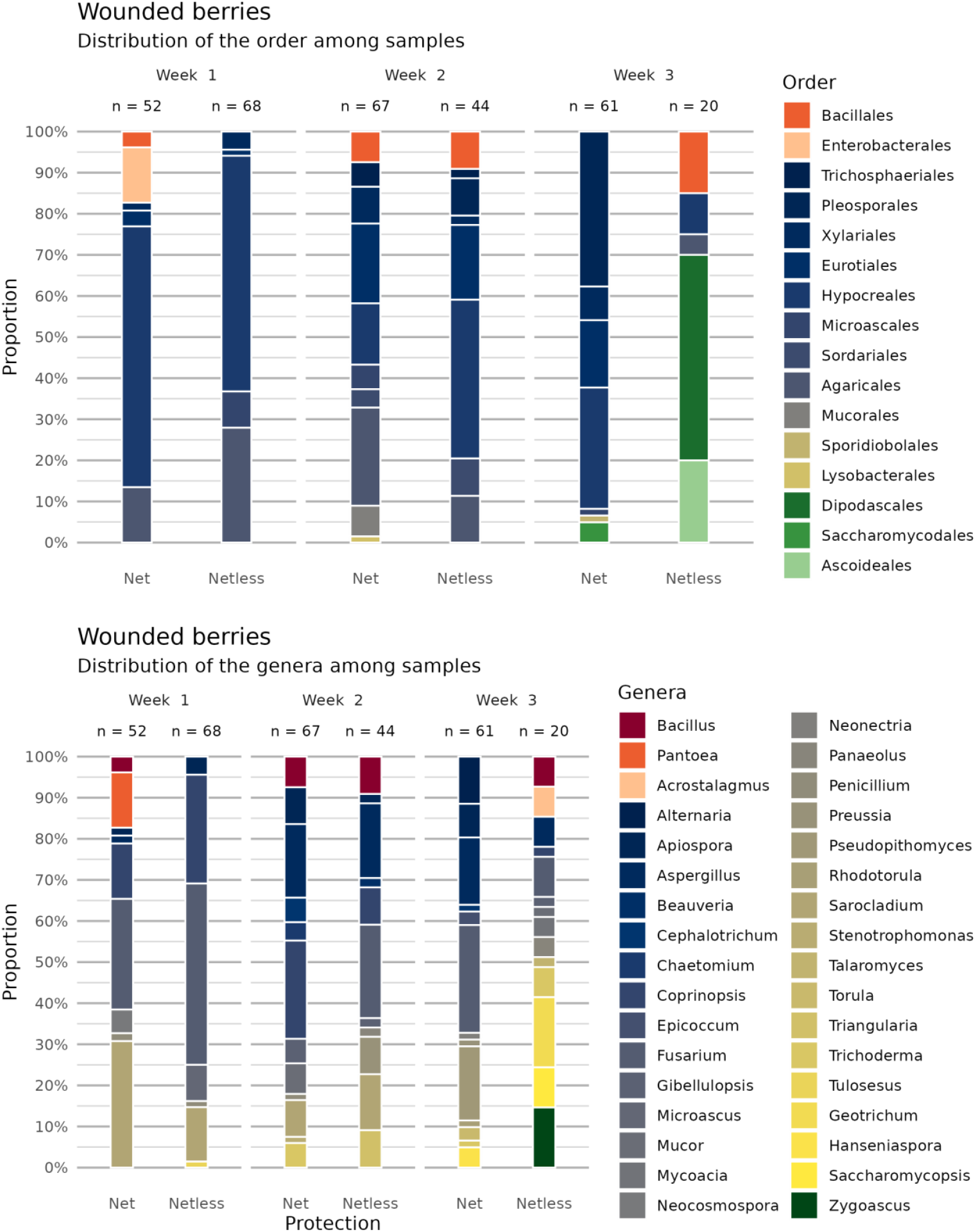
Evolution over time of the microbial community inferred from the relative abundance of fungal and bacterial orders and genera in berries of Chasselas grapes sampled from artificially wounded grape clusters protected or not by a net in a vineyard plot at Nyon (see Table S2 for the classification of the isolates).

### Experiment 3: Comparison of the fungal communities associated with sour rot symptomatic berries, with healthy berries (surface-sterilized or not), and on the surface of Drosophila flies

A sour rot infected vineyard of the cultivar Gamay in Begnins was used to compare the fungal community found on and in healthy grapes, in sour rot diseased grapes and on the surface of potential insect vectors. Fungi were isolated from the surface of healthy berries, from the surface of surface sterilized healthy berries, from sour rot affected berries and from the surface of captured drosophilids in the experimental vineyard. A total of 1020 fungal isolates were obtained in pure culture, 29 from surface-sterilized healthy berries, 265 from non-surface-sterilized healthy berries, 507 from berries affected by sour rot and 219 from the surface of *Drosophila* spp. (Table S3, https://github.com/agroscope-ch/sour-rot/). Among these isolates, ITS sequences were obtained from 999 isolates. The 100% similarity assembly performed on the 999 obtained ITS sequences showed that our isolates comprised 136 different ITS genotypes. Further BLAST analyses of these genotypes in GenBank enabled the identification of 40 genotypes at the species rank, 92 at the genus rank, one at the family rank, two at the order rank and one at the class rank. Yet, BLAST analyses gave results that did not allow us to assign two ITS genotypes to an order. The top score sequence for two genotypes (Table S3) were only assigned to a fungal order (*Curvibasidium pallidicorallinum*) or a fungal class (Sordariomycetes sp.) in GenBank. Yet, ITS sequences could not be obtained for one fungal morphotype (21 isolates) and it was consequently designated as “fungal sp.”.

The structure of the fungal communities isolated from pulp and skin of healthy surface-sterilized berries, from the surface of healthy non-surface-sterilized berries, from sour rot symptmatic berries and from the surface of *Drosophila* spp. was first examined at the order rank (Figure 7). Healthy surface-sterilized berries harbored very few fungi, having less than 13.4% of the fungal community in common compared with the communities isolated from the surface of healthy non-surface-sterilized berries or the *Drosophila*, and even less than 0.6% with the community isolated from sour rot-affected berries (Table S3). While the fungal community isolated from the skin and pulp of healthy surface-sterilized berries was dominated by Xylariales and Dothideales with relative abundances of 41.4% and 27.6%, respectively, the community associated with the surface of unsterilized healthy berries was dominated by Pleosporales (32.8%) and Saccharomycotina (34.4%; Serinales (8.7%), Saccharomycodales (18.5%), Pichiales (6.4%) and Pfaffomycetales (0.7%)). With 81.2% and 76.7%, the subphylum Saccharomycotina represented the vast majority of fungi associated with sour rot affected berries and those present on the surface of *Drosophila* spp. respectively, both modalities dominated by the orders Saccharomycodales and Pichiales. Notably, yeasts (Saccharomycotina), typically reported to be associated with sour rot, were completely absent from surface-sterilized berries.To compare the three remaining modalities with each other at genus/genotype level (Figure 8; Table S3, columns B-D), we used datasets consisting of mycobiomes isolated from the surface of healthy berries (*n*=40), berries showing symptoms of sour rot (*n*=66), and the surface of drosophila flies (*n*=18). We compared the relative abundance of ITS genotypes of isolated fungi, identified according to the results of similarity searches in GenBank, between these three datasets. The reason we rely on relative abundance is to be able to compare species distributions in each of the modalities, independent of sampling effort. Our data (Figure 8 and Table S3) indicate that some of the genera found in sour rot symptomatic berries were not shared between the surface of healthy berries and fruit flies. For instance, the genera *Metschnikowia* (Serinales) and *Aureobasidium* were not detected on *Drosophila* spp., while present on the surface of healthy berries and in sour rot affected berries. On the opposite, the genus *Mucor* was absent from the surface of healthy berries but present on flies and in diseased grapes. Looking at sour rot associated fungi, the two most abundant genera were the yeast genera *Hanseniaspora* (Saccharomycodales) and *Pichia* (Pichiales). Fungi from the genus *Hanseniaspora* were isolated from all three modalities, but for example the genotypes *H. uvarum* gen. 8, *H. cf. uvarum* gen 1, gen 4 and gen 6 were more abundant in sour rot affected berries than on healthy non-surface-sterilized berries or drosophilids (see Table S3 and Figure 8). Some *Hanseniaspora* genotypes were exclusively present in sour rot symptomatic berries or only in two out of the three modalities (e.g., *H*. aff. *uvarum* gen 10, *H*. cf. *uvarum* gen 12 and 3). The genus *Pichia* consisted out of 38 ITS genotypes and BLAST top scores identified four different species, namely *P. californica, P. kluyveri* and close allies [cf.], *P. terricola* and close allies [cf.] as well as *P. sporocuriosa*. In addition, two potentially other species (i.e., *P. kudriavzevii* [cf. or aff.], *P*. cf. *fermentans*) were identified along with 12 other unidentified *Pichia* ITS genotypes. From the 38 ITS genotypes obtained for the genus *Pichia*, 18 were only isolated from sour rot affected berries. *Pichia californica* and *P. terricola*, the two most abundant genotypes in sour rot affected berries, were also isolated from the surface of *Drosophila* spp.. Four other *Pichia* genotypes were shared among the three modalities, although three of them were more abundant in sour rot diseased grapes than on healthy berries or on flies (e.g., *P*. cf. *terricola* gen 1, *P*. cf. *kluyveri* gen 2, and *P. kluyveri*). Among the five ITS genotypes of *P. kudriazewii* and allies, only one was shared among the three modalities (i.e. *P*. cf. gen. 1; Table S3), but with a lower abundance in sour rot affected berries than on healthy non surface-sterilized berries or on *Drosophila* spp.. The remaining four genotypes were isolated exclusively from the surface of healthy berries (cf. gen. 3), from the surface of *Drosophila* spp. (cf. gen. 2 and 4) or from sour rot affected berries (aff.).

**Figure 7.**
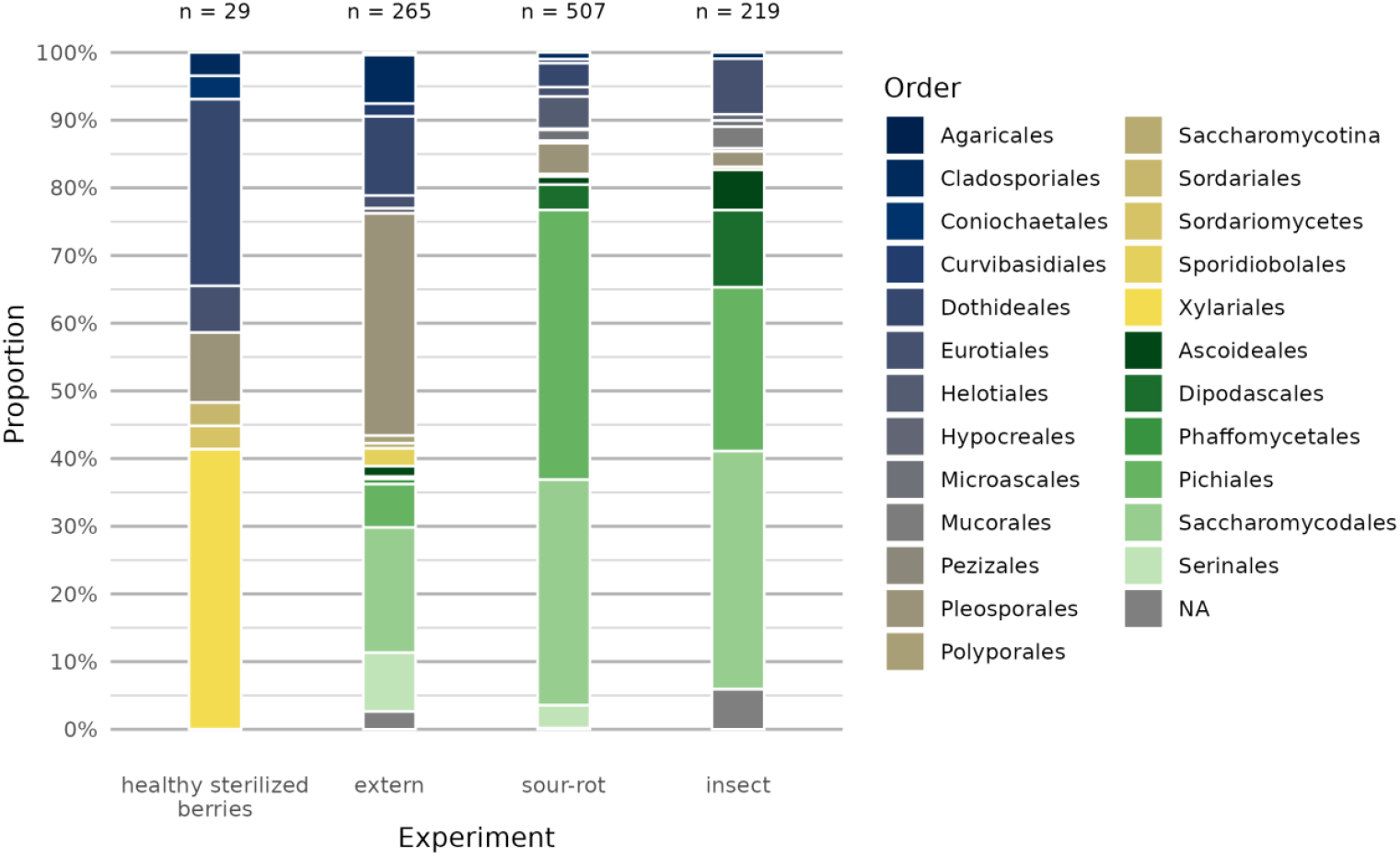
Relative abundance of fungal orders, where this classification rank was indicated in GenBank, in healthy berries post-surface sterilization (internal), on the surface of healthy non-sterilized berries (external), in berries affected by sour rot (sour rot) of grapes cv. Gamay, and on the surface of *Drosophila* spp. (insects) captured in a commercial vineyard in Begnins.

**Figure 8.**
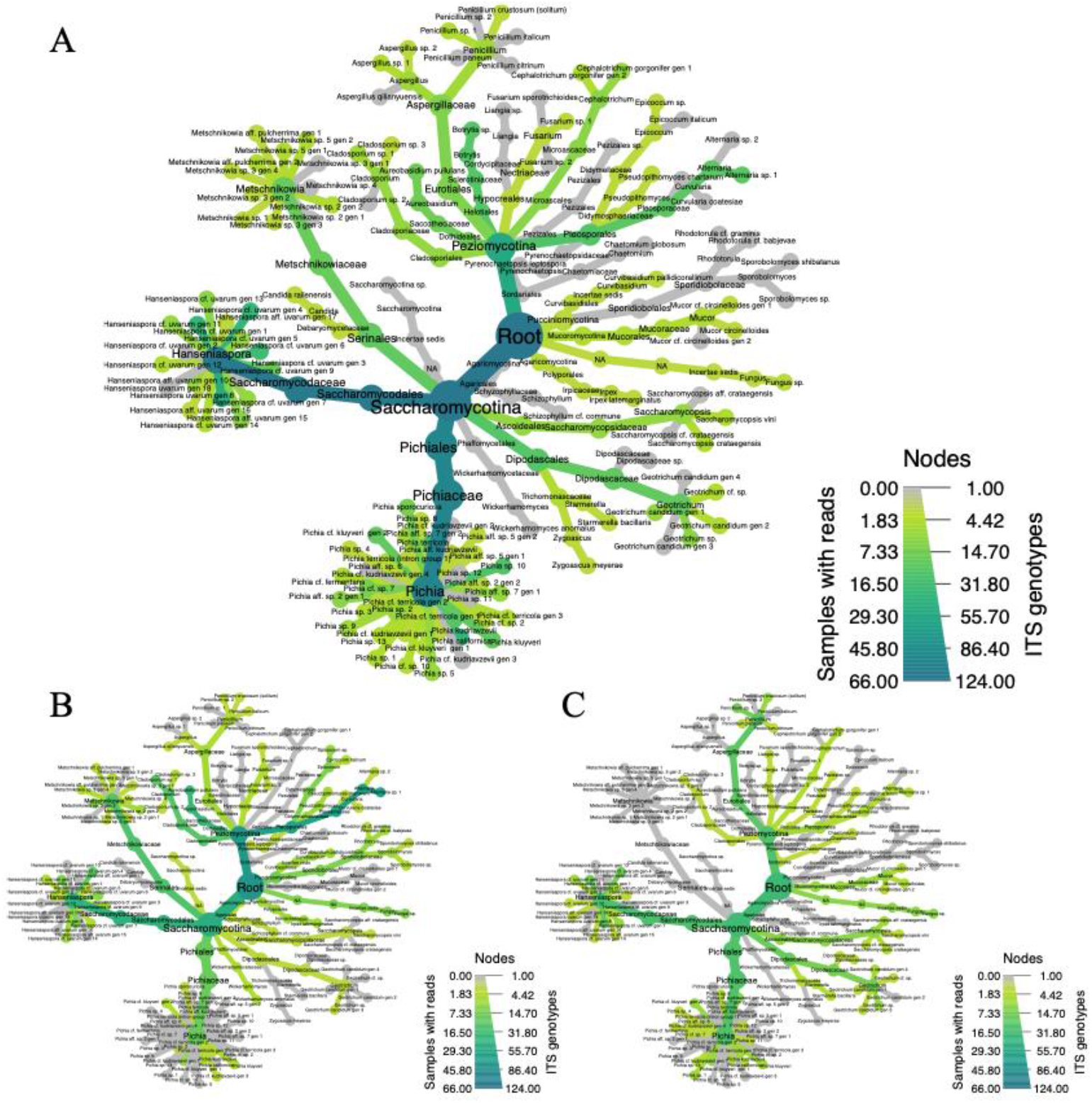
Abundance (in counts) of the fungal ITS genotypes in the three modalities (A. sour rot symptomatic berries, B. surface of healthy berries, C. surface of *Drosophila* spp.). The width of the edges denotes the number of counts in the whole experiment, while the color indicates the number of counts in the considered modality. The color scale is different for each group. The counts are not normalized, i.e. not proportional.

In addition, we isolated six other genera of Saccharomycotina, namely *Candida, Geotrichum, Metschnikowia, Saccharomycopsis, Starmerella, Zygoascu*s as well as an unidentified Dipodascaceae. Two Isolates of *Candida railenensis* were present in sour rot affected berries. Representatives of *Geotrichum* (Dipodascaceae) were mostly associated with sour rot diseased berries but they were also isolated from the flies and from the surface of healthy berries. Representatives of *Metschnikowia*, completely absent from the surface of *Drosophila* spp., were more abundant on the surface of healthy berries than in sour rot affected berries with half of the genotypes shared between the two modalities. Members of the genus *Saccharomycopsis* were more abundant on the surface of drosophilids than in the other two modalities. *Zygoascus* and the unidentified Dipodascaceae were each represented by a single isolate present in sour rot affected berries and on the surface of healthy berries respectively. Four isolates of *Starmerella* were identified on drosophilids and one in sour rot affected berries.

Two additional species commonly considered as pathogens were also isolated (Table S3, Figure 8). One of these was a *Botrytis* species, which was exclusively present in berries affected by sour rot. Yet, our ITS sequences could not be assigned at the species rank, as four *Botrytis* species have identical ITS sequences in GenBank, namely *B. cinerea, B. eucalypti, B. fabae* and *B. pelargonii*. Also, an unidentified *Alternaria* species (sp. 1) was particularly abundant on the surface of healthy berries, with an ITS sequence 100% identical to type sequences deposited in GenBank for three different species (i.e., *A. alstromeriae, A. cerealis* and *A. aborescens*) and to several sequences deposited under the name *A. alternata*. Yet for the latter species, no type sequence is flagged in GenBank.

Pairwise comparisons of the relative abundance of the fungal ITS genotypes between the three mycobiomes also indicated that genus *Alternaria* was more abundant on the surface of healthy berries than in the two other modalities (Figure 9). The Aspergillaceae family, mainly genus *Penicillium*, as well as the order Dipodascales were inferred to be essentially associated with the surface of insects, while the genus *Hanseniaspora* was found to be more abundant in sour rot symptomatic berries than in the two other modalities, except for *H. uvarum* genotype 8, which was more abundant on the surface of drosophilids than on the surface of healthy berries. Genus *Pichia* appeared more abundant on insects’ surface and in sour rot symptomatic berries than on the surface of healthy berries. Overall, these three pairwise comparisons indicate that the mycobiome of sour rot symptomatic berries is somehow closer to the one isolated from the surface of the insects than to the one isolated on healthy berries.

**Figure 9.**
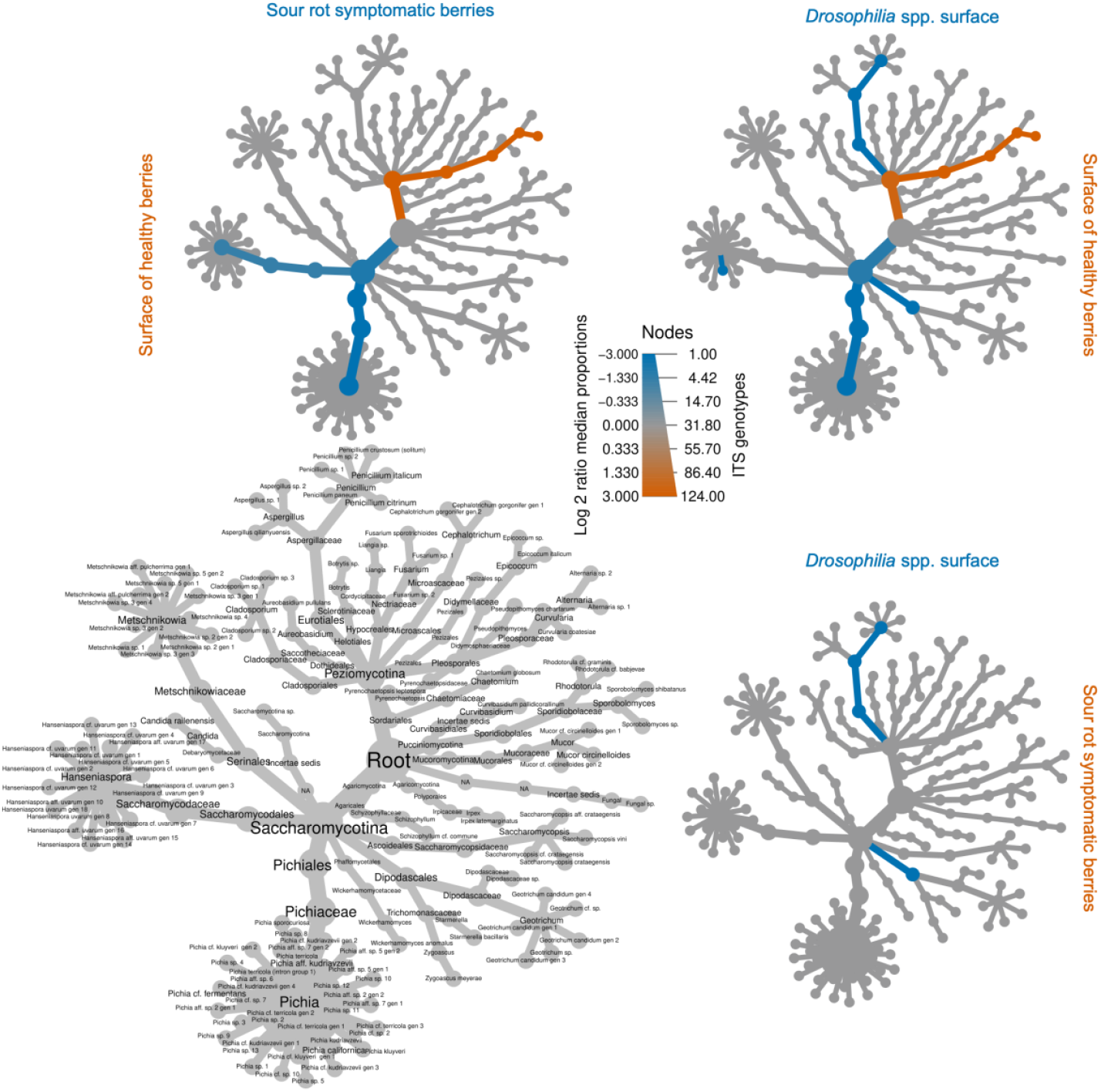
Pairwise differences of log (in base 2) median relative abundance of ITS genotypes in the three groups (healthy berries, sour rot symptomatic berries and *Drosophilia*). For each edge (i.e. taxon), the median of the proportion of the corresponding taxon in the modality considered is computed. Then, the log (in base 2) of the ratio of the two medians (or equivalently, of the differences of the logs) is represented on the color scale. Note that, by convention, if any of the two medians is 0, then 0 is displayed. See this [hyperlink](https://grunwaldlab.github.io/metacoder_documentation/faq.html#differential-heat-trees-with-compare_groups-which-color-is-which-treatment) for implementation details and further clarifications. A taxon colored in blue is more abundant in the modality in the column while a taxon colored in orange is more abundant in the modality of the row (see the color of the labels of rows and columns).

Finally, we sought to determine whether the introduced *D. suzukii* carried the same or different sour rot associated yeasts as endemic *Drosophila* species. Our data show that the surfaces of endemic vinegar flies and *D. suzukii* (Table S3; Figure 10) carry a similar load of yeasts, 81.0% versus 80.6% of the fungal isolates, respectively. On the surface of all *Drosophila* species the genus *Hanseniaspora* was the most abundant with 33.3% for endemic species and 36.4% for *D. suzukii*, followed by the genus *Pichia* with 31.1% and 19.4% for the native and introduced species, respectively. These two genera accounted for over 55% of the yeasts present on the surface of introduced and endemic *Drosophila* species. The other Saccharomycotina genera present on the surface of *D. suzukii* were *Geotrichum* (12.4%) and *Saccharomycopsis* (3.1%), two genera also present on the surface of endemic species (4.4% and 10.0%), while the genus *Starmerella* (3.1%) was only present on the surface of *D. suzukii*, and the representative of *Dipodascaceae* only isolated from endemic *Drosophila* species (1.1%).

**Figure 10.**
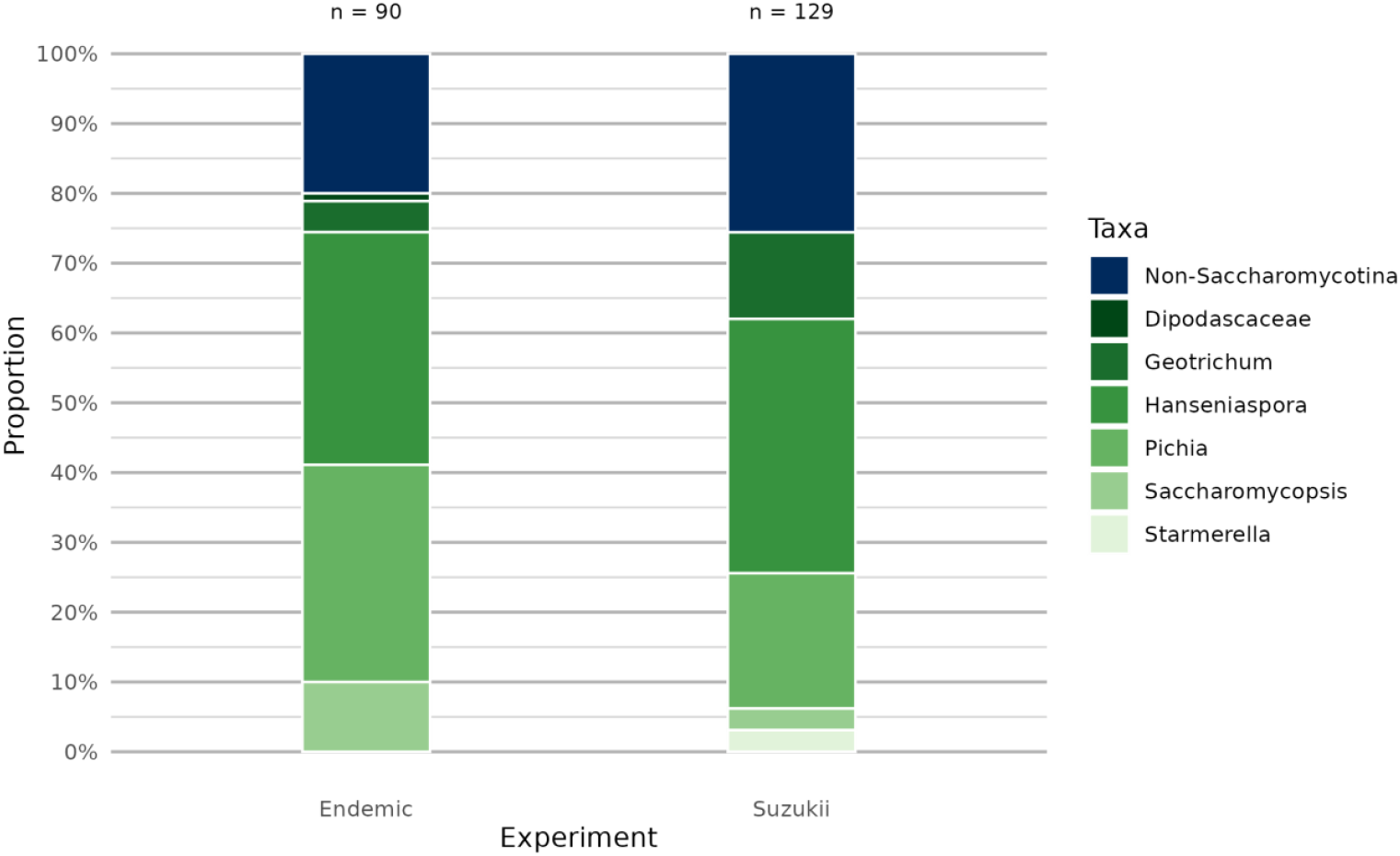
Comparison of Saccharomycotina genera present on the surface of *D. suzukii* and on the surface of endemic *Drosophila* species. Fungal genera not associated with sour rout were coded as non-Saccharomycotina.

## DISCUSSION

### Sequence-based identification of the isolated microorganisms

In this study, we assigned names to the isolated microorganisms based on the results of sequence similarity searches (BLAST top score sequences) in GenBank for ITS, the fungal barcode locus (Schoch et al. 2012) and for the 16S, the bacterial barcode locus (Lebonah et al. 2014). By using a molecular based identification of these organisms and adopting a 100% sequence similarity threshold, we were able to identify less than one third of the fungi and half of the bacteria at the species rank (Tables S1-S3). The 16S locus has been shown to offer resolution at genus rank for bacteria but in most cases not below (Bartoš et al. 2024). The ITS locus is known to fail in the recognition and delimitation of cryptic species in many Ascomycota genera. As examples (see Tables S1-S3), identical ITS sequences have been deposited in GenBank under different species names and this locus offers very low identification power for the discrimination of cryptic species in genera such as *Alternaria* (Dettman & Eggertson 2022), *Botrytis* (Plesken et al. 2021), *Cladosporium* (Schubert et al. 2007), or *Fusarium* (Crous et al. 2021). To unambiguously identify these cryptic species, a multi-locus phylogenetic analysis using a representative sampling of species in each genus will be necessary. The poor resolution offered by ITS for the discrimination between cryptic mold species also applies to yeasts, the order Saccharomycotina representing more than 95% of the fungi isolated in this study.

About ten years ago, Saccharomycotina (Groenewald et al. 2023; previously Saccharomycetales) consisted in 70 genera (Kurtzman & Robnett 2013) versus circa 100 genera today according to the GenBank taxonomy (https://www.ncbi.nlm.nih.gov/Taxonomy). However, the sequences deposited in GenBank for the most frequently isolated genera in the present study (i.e., *Hanseniaspora, Metschnikowia*, and *Pichia*), have often been deposited with the suffixes cf., aff. or sp. and we therefore restricted ourselves to this nomenclature. The genus *Candida* has long been the taxonomic dustbin of yeasts, resulting in many unclassified sequence deposits for this genus in GenBank (see Table S3). Nomenclatural problems with yeast species were already highlighted by Douglass et al. (2018) who assessed, based on genome sequencing, that *Candida krusei, Pichia kudriavzevii, Issatchenkia orientalis* and *Candida glycerinogenes* were in fact all the same species. Also, *Candida californica* was recently transferred to the genus *Pichia* (*P. californica*; Groenewald et al. 2023). As Saccharomycotina have small genomes, it has been suggested that genome sequencing is cheaper and that it will be easier to identify cryptic and closely related species in this order in comparison to obtain sequences for multi-locus phylogenetic analyses (Koufopanou et al. 2013). Genome sequencing has successfully be used for the large-spored *Metschnikowia* clade (Lachance et al. 2016). At present, a multigene or genomic phylogenetic approach is however not applicable to the study of fungal communities that involve hundreds of isolates. A second limitation comes from the possibility of intra-genomic variability in the ribosomal tandem repeats as indicated by Colabella et al. (2018). In their study, authors obtained from single fungal isolates of the genus *Candida* several different ITS. Yet, such a variability in the ITS of a single strain has, to our knowledge, not been reported so far for other Saccharomycotina.

Due to these practical limitations, we refrained from directly adopting the species names associated with the BLAST top score for our ITS sequence genotypes for Saccharomycotina. This decision was made even though the results assigned several different ITS genotypes to the same species (i.e., *Hanseniaspora uvarum, Pichia terricola*, etc.). We therefore often adopted the suffixes cf. (probably that species) or aff. (close but could be another species) rather than taking the risk to misidentify our isolates (Table S1-S3; Hofstetter et al. 2019). The only Saccharomycotina for which we adopted a species name associated with BLAST top score was when a single ITS sequence obtained from a type specimen showed 100% of similarity to our isolate. To clarify the taxonomy of Saccharomycotina and delimit species within the genera of this fungal order, multi-locus or genomic phylogenetic analyses as well as ITS sequences for type collections are imperatively needed.

### Microbial communities associated with sour rot and grey mould symptomatic grape clusters

A comparison of the microbial communities in grape clusters symptomatic for sour rot or grey mould within the same vineyard plot revealed clear distinctions between net-protected clusters affected by grey mould and unprotected clusters affected by sour rot (Figure 1). Although the two fungal genera *Penicillium* and *Pichia* were associated with both diseases, the remaining ten fungal genera were only isolated from clusters affected by one or the other disease. Sour rot symptomatic clusters predominantly hosted yeast genera from the order Saccharomycotina (i.e., *Candida, Geotrichum, Pichia*, and *Saccharomycopsis*) as well as the bacterial genus *Acetobacter*, confirming thereby the results of previous studies (Bisiach et al. 1982; Guerzoni and Marchetti 1987; Barata et al. 2012b; Hall et al. 2019a; Gao et al. 2020). However, Saccharomycotina were much less abundant in grey mould than in sour rot symptomatic clusters. Grey mould symptomatic clusters hosted principally non-yeast genera such as a *Botrytis* species of the *B. cinerea* clade and four other fungal genera absent in sour rot affected clusters. All the species in the *B. cinerea* clade are considered as grapevine grey mould agents. Overall, only the bacterial genus *Pantoea* was isolated from grey mould affected clusters, but no acetic acid bacteria (AAB). Despite our limited sampling efforts, the observed differences between the microorganisms associated with sour rot and grey mould confirm that they are two distinct diseases as suggested by Crandall et al. (2022).

We observed in our Chasselat vineyard plot that grey mould developed only on clusters protected by a net, while sour rot developed only on clusters that remained unprotected. These results are consistent with previous studies and emphasize that when insect vectors such as vinegar flies are present and have access to the grapes, sour rot predominates, but when potential vectors are lacking (as in the case of net-protected clusters), grey mould is able to develop. Yet, in our third experiment we were nonetheless able to identify the presence of a *Botrytis* sp. in relatively high abundance in sour rot affected grape clusters, while this genus was absent on either the surface of healthy berries or on drosophilids (Figure 8; Table S3). The ITS BLAST top score genotype for this *Botrytis* sp. was 100% similar to the sequences of four different *Botrytis* species, all belonging to the *B. cinerea* clade. Although our initial observations (Figure 1, Table S1) suggest that sour rot supplants grey mould when both diseases are present in a vineyard (Bisiach et al. 1982; Hall et al. 2018), the outcome of the interaction may vary, as *Botrytis* is also capable of developing in grapes affected by sour rot (Figure 8). This result suggests that grey mould may coexist with sour rot and that it is not being systematically stopped by sour rot at an advanced stage of ripening as suggested by Bisiach et al. (1982).

### Sour rot associated microorganisms

Comparing the microorganisms isolated from sour rot symptomatic grape clusters (Figures 1B, 2, 6, 8; Tables S1-S3), we identified the same sour rot disease-associated Saccharomycotina compared to previous studies (Barata et al. 2008, 2012a; Hall et al. 2018, 2019a; Gao et al. 2020). The fungal community of sour rot affected grapes is essentially composed of the yeast genera *Hanseniaspora, Pichia* (previously *Issatchenkia*), *Saccharomycopsis*, and *Zygoascus*. We also report for the first time yeasts of the genus *Geotrichum* to be associated with grape sour rot. The yeast genus *Geotrichum* was isolated from sour rot symptomatic clusters in all three experiments (Figures 2, 6, 8; Tables S1-S3). Species of that genus have recently been identified as the principal agents for post-harvest sour rot in different fruits and vegetables such as jocote (*Spondias purpurea*), tomato or potatoes in Brazil (Paes et al. 2021). Additionally, Zhu et al. (2016) showed that yeasts of the genus *Geotrichum* are able to metabolize sugars in acetic acid. Similarly, it has been reported that the yeast *Hanseniaspora uvarum* can produce oxidation defects in wine by increasing acetic acid, ethyl acetate and acetaldehyde content (Clemente-Jimenez et al. 2004; Garde-Cerdan & Ancín-Azpilicueta 2006). Finally, results from a recent study (Lozano et al. 2024) indicate that species of the genus *Candida* and *Hanseniaspora* are able to transform sugar into acetic acid. Together with the aforementioned studies, our results support the hypothesis that fungi can induce sour rot even in the absence of bacteria.

Representatives of the bacterial genus *Acetobacter* were isolated only in our first experiment and only at one location, in the canton of Vaud; Figure 1). Hall et al. (2019a) found *Acetobacter aceti* and *A. pasteurianis* in sour rot symptomatic berries, but not the two species isolated in the present study (i.e., *A. fabarum* and *A. gahnensis*; Table S1). In the present study, clusters were tested for their acetic acid content to determine if they were really affected by sour rot using an acetic acid threshold above 0.83 g/L as defined by Hall et al. (2019). Although AAB were absent in some symptomatic clusters from the Grisons (Figure 1) and also in the second experiment (Figure 3-5), the juice from these grapes exceeded the acidity threshold, and the clusters were therefore considered affected by sour rot. In our second experiment, only bacteria from the genus *Bacillus* (Bacillales) were present in sour rot affected clusters, a genus also isolated from diseased grapes in previous studies but reported not to be implicated in the expression of sour rot (Gao et al. 2020, Hall et al. 2019a). In fact, we isolated AAB only in our first experiment where clusters were left for two weeks for microorganisms’ enrichment at room temperature before culturing the microorganisms, which evidently accentuated the ripening stage of the berries. In the other two experiments, we did not enrich the microorganisms and processed the bunches immediately after collection. Nonetheless, if present, we should have been able to isolate AAB from sour rot–affected berries in the second experiment, particularly given the considerably greater sampling effort compared to the first experiment and the proven suitability of our methodology for isolating and sequencing *Acetobacter* species (Figure 1). Our results therefore suggest that AAB may appear late in the development of sour rot and that fungi might be able to initiate the catalyzation of acetic acid by themselves, even in the absence of AAB.

However, these results should be viewed with caution as we used a single medium to grow all microorganisms, which probably led to an underestimation of the diversity and proportion of bacteria present in our grape samples. While PDA is considered the best culture medium for fungi (Devi et al. 2018; Black 2020), the study of bacterial communities requires specific and/or multiple culture media in order to maximize their biodiversity (Bonnet et al. 2019; Lagier et al. 2015). Studies using high-throughput sequencing of bunches showing symptoms of sour rot (Gao et al. 2020; Hall et al. 2019a) obtained higher bacterial abundance and diversity than we did, as well as a higher proportion of *Acetobacter* and *Gluconobacter*, two genera that were absent in our second experiment (Figure 3). Barata et al. (2012) reported that AAB (*Acetobacter* and *Gluconobacter*) were abundant as soon as bunches showed symptoms of sour rot. In addition, the study by Hall et al. (2019a) showed that the bacterial community in grapes expressing sour rot symptoms was very different from place to place, with, for example, the genus *Gluconobacter* and the order Bacillales very rarely present in disease-affected bunches in New York State (USA), while very abundant in Tasmania (Australia). In our second experiment, we isolated only Bacillales, an order also abundant in the study by Gao et al. (2020) but absent in the study by Barata et al. (2012). Consequently, the abundance and diversity of AAB appear to depend on the stage of ripening of the grapes and the geographic locality where they are sampled. Our second experiment took place over the three weeks prior to harvest and in a single vineyard plot. Sampling the bunches in a single plot could, at least in part, explain why bacterial diversity was low. The stage of rotting of the bunches, even though four bunches had reached the significant level of acetic acid characterizing sour rot, could also be responsible for the failure to isolate AAB in our second experiment. As suggested by Barata et al. (2012) for lactic acid bacteria, and by our first experiment for AAB, an enrichment step may have increased the bacterial diversity and the abundance of AAB. A two weeks prolongation of our sampling period in our second experiment might therefore have favored the isolation of AAB.

Further research is therefore needed to confirm that the genera *Acetobacter* and *Gluconobacter*, reported to cause sour rot in Europe (Barata et al. 2012), North America (Hall et al. 2019a) and China (Gao et al. 2020), may not be obligatory for disease expression. Besides, further studies might also verify that that yeasts of the *Geotrichum, Candida* and *Hanseniaspora* genera are capable of initiating sour rot expression in the absence of AAB will also require further study.

### Determining the impact of wounds and insect vectors on the microbial community

When comparing the microbial communities of adjacent healthy berries with those affected by the disease, and using berries taken from disease-free grape clusters as controls (Figure 3), our data suggest that sour rot disease is not directly spreading from symptomatic berries to adjacent healthy berries. Sour rot affected grapes are characterized by a high relative abundance of yeasts (Saccharomycotina, 70%) and the presence of bacteria (Bacillales), that characterize berries affected by sour rot, while asymptomatic berries hosted only few Saccharomycotina and no bacteria. This observation is consistent with previous studies (Barata et al. 2012b, Gao et al. 2020, Hall et al. 2019a) and suggests that insect vectors such as drosophilids play an important role in the dissemination of the disease.

Regardless of insect access to the grapes, the microbiomes on the surface of intact berries were relatively similar three weeks after the start of the second experiment (Figure 4). None of the grape clusters sampled in this experiment reached the acetic acid threshold associated with sour rot. This reflects probably the relatively low abundance of Saccharomycotina (i.e., < 20%) on healthy grapes, whether vinegar flies had access to the berries or not. It therefore seems that the presence of insect vectors had not a major effect on the evolution of the grape microbiome in the absence of skin wounds. Wounds were identified here as the most important factor promoting the development of sour rot. Wounds are generally considered as necessary for disease development as also stated by McFadden-Smith and Gubler (2015). Rainy weather at an early ripening stage (Vogel et al. 2021), together with warm temperatures and a sugar content in grape berries above 15 °Brix, has been identified as the main factors leading to sour rot expression in North America (Hall et al. 2018).

Although wounding was identified as the main factor in the expression of sour rot in our second experiment, the presence of insect vectors also appears to play a key role in the onset of symptoms, accelerating the evolution of the microbiome towards a high proportion of Saccharomycotina in wounded berries compared with unwounded berries by around a week.(Figure 5). Cracks in the grape skin enable microorganisms to access nutrients in the grape pulp, thereby facilitating the development of Ascomycota and in particular yeasts. Consequently, the role of vinegar flies in the expression of sour rot is suggested to be primarily mechanical (Barata et al. 2012b, Hall et al. 2018). According to Kehrli and Linder (2018), the activity of hatched *D. suzukii* larvae in grape berries delays the healing of their skin by around two weeks, a delay also observed by Barata et al. 2012a or Rombaut et al. 2017, while the wounds around older, dead eggs are completely healed. Our results are consistent with these previous studies, suggesting that wounds heal more slowly when insects are attracted to the wounds, allowing fungi and bacteria to access the berry pulp and to initiate a microbial community that leads to sour rot expression. Our results also support the observations of Hall et al. (2019a) indicating that sour rot-associated yeasts are present on the surface of intact berries but cannot reach the pulp without skin wounds. This is further evidenced by the similar evolution of microbial communities in wounded berries at both the order and genus levels, regardless of vinegar fly access (Figure 6A and 6B; Table S2). Our data, however, emphasize that insect vectors only started to influence the microbiome composition in wounded berries after three weeks, particularly by boosting yeast development (Saccharomycotina, genera *Geotrichum, Saccharomycopsis* and *Zygoascus* – see Table S2), and this while net-protected wounded berries predominantly hosted Pleosporales species. Nevertheless, a few Saccharomycotina were also isolated from net-protected wounded berries, suggesting that sour rot might develop later, even in the absence of insect vectors such as drosophilids. To verify this hypothesis, future studies should monitor the evolution of the microbiomes in wounded berries protected by a net over a longer period.

### Does sour rot expression result from the microbiome on the surface of healthy berries and/or from the microbiome on the surface of drosophilids?

The microbiome associated with sour rot symptomatic berries is presumed to originate either from the penetration of surface-resident fungi and bacteria or through contact with the microbiome carried by visiting insects (Hall et al. 2019a). Here, we did not consider the transport of spores in the air because a recent study indicated that the airborne microbiome has a minor contribution to the grape microbiome (Abdelfattah et al. 2019).

The number of fungi isolated from surface-sterilized berries was ten to seventeen times lower than on unsterilized or symptomatic berries. Such differences may be partially explained by a larger number of berries sampled from sour rot affected clusters (64 berries/507 fungal isolates; Table S3) but this does explain the observed differences between surface-sterilized berries and unsterilized berries as the same numbers of berries were sampled (50 berries for both conditions and 29 versus 265 fungal isolates obtained respectively). The mycobiome of surfaced sterilized berries consisted mainly of species belonging to the order Xylariales (particularly the genera *Annulohypoxylon*,

*Biscogniauxia, Daldinia* or *Nemania*; see Figure 7; Table S3). Many Xylariales species are known to be common plant endophytes (Becker and Stadler 2021; Ma et al. 2022, U’Ren et al. 2016). Yet, the surface-sterilized berries did not only host fungi from different systematic groups compared to those from the other three modalities, but no yeasts were isolated from these surfaced sterilized berries (Figure 7; Table S3). Our results thus contradict those reported by Hall et al. (2019b), who isolated mainly yeasts from surface-sterilized healthy berries. However, we think that this might be explained by the different condition of the analyzed grapes. Indeed, Hall et al. (2019b) sampled grapes in vineyards one to five days before harvest, but also table grapes obtained from supermarkets, where these were most likely kept in cold storage several days before being sold. It is conceivable that microfissures developed in the grapes on sale, allowing surface-resident yeasts on commercialized and late-harvested berries to penetrate the fruit. While the mycobiome present on the surface of healthy grapes has been widely studied, our study, as well as the study by Hall et al. (2019b), are to our knowledge, the only studies that investigated the fungi living within the pulp of healthy grape berries. Further research is needed to confirm whether Saccharomycotina can thrive as endophytes of healthy berries or if they are only present on the surface of grape berries as suggested by our data.

Our results showed that, at the order level, the fungal community associated with sour rot symptomatic berries was more similar to that isolated from the surface of vinegar flies than to the community isolated from healthy berries (Figure 7; Table S3). This was particularly evident for yeasts, which constituted the vast majority of fungi associated with symptomatic berries but also of those isolated from the surface of *Drosophila* adults, and this when yeasts represented less than a third of the fungal community isolated from the surface of healthy berries. Consistent with previous observations, fruit flies appear to enhance the presence of yeasts in sour rot symptomatic berries, particularly those of the genera *Hanseniaspora* and *Pichia* (Table S3; Figure 8). This finding is reinforced by the pairwise comparisons of the relative abundance of these two genera being more frequent in the mycobiomes associated with sour rot affected berries and on the surface of vinegar flies compared to the surface of healthy berries (Figure 9). A high intraspecific diversity of *H. uvarum* has been documented in winery environments (Grangeteau et al. 2015), as well as an intraspecific nucleotide diversity based on multi-locus typing analyses (Saubin et al. 2020). We also retrieved many different ITS genotypes for *Hanseniaspora*, all highly similar to sequences deposited as *H. uvarum* in GenBank (Table S3; Figure 8-9), but with certain genotypes being more prevalent in some of the experimental setups (i.e., gen 8 on *Drosophila* surface, Figure 9). Given that *H. uvarum* related sequences corresponded to the most abundant fungi in the third experiment across all three modalities, our results align with previous studies (Barata et al. 2012b; Hall et al. 2018). This species has extensively been identified as one of the most common yeasts in grape musts across various countries (Albertin et al. 2016, Romano et al. 2019) as well as in sour rot symptomatic berries in Europe (Barata et al. 2012a). The most prevalent ITS genotypes of *Pichia* were shared among the surface of *Drosophila* spp., sour rot symptomatic berries and the surface of healthy berries, except for *P. californica* absent on the surface of the latter (Figure 8; Table S3). However, not all *Pichia* species appear to proliferate equally in diseased grapes. *Pichia californica, P. kluyveri* et *P. terricola* (including closely related species cf. or aff.) accounted for over half of the *Pichia* strains isolated from symptomatic berries, with the most abundant genotypes being more prevalent in sour rot symptomatic berries than on the surface of *Drosophila* flies or on healthy grape berries (Table S3; Figure 8). These three *Pichia* species therefore seem to have a greater impact in the process leading to sour rot than the other *Pichia* spp. Similarly, Hall et al. (2019a) observed that *P. kluyveri* was the most common species in sour rot symptomatic berries. On the other hand, half of the six ITS genotypes for *P. kudriazewii* were exclusively isolated from the surface of healthy berries and/or drosophilids, while two other genotypes were equally or less abundant in sour rot affected berries compared to the other two modalities (Table 3; Figure 9)). Also, according to Fleet (2011), *P. kudriavzevii* can be found in the soil and on the outside of fruit and vegetables, often in the presence of other *Pichia* species, but is not considered a species responsible for food spoilage, which is in line with our results. It therefore seems that *P. kudriazewii* is less adapted than *P. kluyveri* and *P. terricola* to survive and proliferate in ripening grapes. The Genera *Geotrichum* and *Saccharomycopsis* were predominantly isolated from sour rot affected berries and from drosophilids. Vinegar flies thus seem to contribute more of these two genera to sour rot affected grapes than healthy berries. On the other hand, the genus *Metschnikowia* was absent on the surface of *Drosophila* species but relatively abundant in symptomatic berries and on the surface of healthy berries. It might therefore be that this yeast genus is not well adapted to be transported by vinegar flies, as yeast adhesion has been demonstrated to be dependent on both the yeast strain as well as on the surface it interacts with (Pawlikowska et al. 2019; Whitehead et al. 2021), an aspect that is certainly worth to be investigated in more detail. Overall, yeasts present on the surface of vinegar flies and of healthy berries seem to contribute to the fungal community associated with sour rot in grapes. However, their contribution does not necessarily involve the same genera, species, genotypes and/or in the same proportions.

### Comparison of the mycobiomes associated with endemic and introduced drosophilids

A mutualistic relationship between *H. uvarum* and *D. suzukii* has been demonstrated in immature raspberries where this yeast seems to promote the development of insect larvae (Chakraborty et al. 2022). There is also evidence that *H. uvarum* is one of the most frequent yeasts in the gut of several *Drosophila* species, including *D. melanogaster* (Chandler et al. 2012). When comparing our fungal isolates obtained from the introduced *D. suzukii* compared to native *Drosophila* species (Figure 10), we conclude that fungal communities among *Drosophila* species are overall quite similar and that all vinegar flies carry a high and equal amount of yeasts with *Hanseniaspora* (*H. uvarum*) being the most abundant genus, followed by *Pichia* (several species). It was suggested that *Pichia kluyveri* and *P. terricola* maintain the same positive mutualistic association with *D. suzukii* on the basis of their abundance in the gut of this vinegar fly (Hamby et al. 2012). Stamps et al. (2012) showed that this relationship also exists for *D. melanogaster*, with the yeast species involved in this mutualistic association including not only *Pichia kluyveri*, but also *Candida californica* (now *Pichia californica*; Groenewald et al. 2023) and *C. zemplinina*. Our results confirm previous studies, with *H. uvarum* and its allies, as well as *Pichia spp*. being also frequent on the surface of both endemic and introduced vinegar flies (Figure 10). As observed by Stamps et al. (2012), *Pichia californica* was isolated only on *D. melanogaster*, while this species was absent on the surface of *D. suzukii*. Overall, all *Drosophila* species seem to play a similar role in the transport and transmission of yeast cells between grape berries and they may thus all contribute to the incidence of Saccharomycotina in sour rot affected berries as suggested by Entling & Hoffman (2020). Given that our study is the first that investigated the contribution of the surface of *Drosophila* species to the fungal community associated with sour rot in grapes, it will be interesting to see if future studies are able to confirm our findings.

## DECLARATION

### Data availability

All sequence data have been deposited in GenBank with corresponding accession numbers. The supplementary material is accessible through the hyperlink https://github.com/agroscope-ch/sour-rot/. In particular the data can be downloaded from the Data folder. All the figures displayed in this paper are provided in an interactive .html format (facilitating inspection) and the scripts that have been used to produce all the figures are also provided with detailed comments (including about preprocessing steps). Some additional plots and analyses that have been not included in the paper can also be found there.

### Authors contributions

SH, PK, KG, J-LW and VH conceived the study; SH and VH collected the data, MW, SH and VH analyzed the data; SH, PK, BB, and VH led the writing of the manuscript. All authors commented on the manuscript.

## ACKNOWLEDGEMENTS

We thank Frédéric Vuichard, Nicole Lecoultre, Eric Remolif, Clara Chevalley, Françoise Klötzli-Estermann and Anne-Laure Fragnière for their technical support.

